# AC electro-osmosis in bacterial biofilms: a cautionary tale for electrophysiology experiments

**DOI:** 10.1101/2025.01.22.634266

**Authors:** Victor Carneiro da Cunha Martorelli, Emmanuel Akabuogu, Raveen Tank, Rok Krašovec, Ian S. Roberts, Thomas A. Waigh

## Abstract

Synthetic cationic fluorophores are used widely as probes to measure the membrane potentials of bacterial cells, eukaryotic cells and organelles, such as mitochondria. An external oscillating electric field was applied to *Escherichia coli* cells using microelectrodes and AC electro-osmosis was observed for the fluorophores, independent of the electrophysiology of the bacteria, giving rise to phantom action potentials. The fluorophores migrate around the microfluidic device in vortices modulating their concentration having decreases or dips in fluorescence. We show that the fluorescent dips are universally present when using cationic fluorophores, such as thioflavin-T, propidium iodide, Syto9 and Sytox Green, with or without *E. coli* cells in the inoculum, when stimulated with AC voltages. This is in contrast to the study of Stratford et al (PNAS, 2019) who claim the existence of action potentials. Furthermore, *E. coli* biofilms also demonstrated similar phenomena with dips in the fluorescence. We measured the relaxation times of the fluorophores experiencing AC electro-osmosis, which depended on the biofilm, the cells and the fluorophores used. PI had the smallest relaxation time and Syto9 the highest. Removing the cells resulted in longer relaxation times and introducing biofilm did not significantly change the relaxation times compared with the single cell experiments. Furthermore, fluorescently labelled DNA and fluorescent colloidal beads also demonstrate fluorescent dips through AC electro-osmosis, showing that these particles can be driven through biofilms. This is the first study of AC electro-osmosis in bacterial biofilms, indicating a surprisingly high mobility of charged molecules within the extracellular polymeric substance, which could be used to treat biofilms i.e. to increase the kinetics of delivery of antibiotics.

## I. INTRODUCTION

Bacterial biofilms are surface adsorbed communities of bacterial cells that are coated in extracellular polymeric substance (EPS). Individual bacteria communicate with each other via quorum sensing to decide whether a surface is suitable to form a biofilm by releasing quorum auto-inducer molecules [1]. Biofilms play an important role in resisting antibiotics and bacteria in biofilms are often tolerant to 2-3 orders of magnitude higher concentrations than individual planktonic cells [2]. Biofilm formation occurs with the majority of bacteria on the Earth [3], with an estimated 5 × 10^30^ bacteria and 10^12^ species [4] [5]. It is thus imperative to study how bacteria communicate to form biofilms to combat infections and, specifically, antimicrobial resistance.

The phenomenon of electrical signalling through small charged ions (such as *K*^+^) has recently been observed, presenting a novel aspect of bacterial communication alongside quorum sensing [6]. This form of signalling enables bacteria to communicate within small clusters and biofilms [7]. Electrical signalling is possible since bacteria and bacterial biofilms are examples of excitable matter, akin to neuronal, sensory, and cardiac cells with voltage-gated ion channels [8] [9]. The pioneering work of Prindle et al. showed *Bacillus subtilis* biofilms that experience nutrient scarcity produce waves of *K*^+^ ions [6]. Similar electrical signalling events have been seen in *Escherichia coli* biofilms subjected to stress induced by exposure to blue light, which generates reactive oxygen species [7]. Hyperpolarization is also observed in mitochondria in response to ROS stress, indicating an evolutionary conserved physiological process [10] [11] [12]. Additionally, wave pulses of hyperpolarization have been observed propagating through biofilms of *Neisseria gonorrhoeae* [13], while *Pseudomonas aeruginosa* has demonstrated hyperpolarization in response to blue light [14]. Spiking phenomena of the membrane potentials of individual cells have also been measured from *E. coli* using optogenetics techniques [15] [16]. These discoveries underscore the diverse ways bacteria employ electrical signalling in different environmental contexts.

Electrophysiology experiments are essential for understanding the intricate workings of biological systems, particularly in neuroscience, cardiology and more recently bacterial cells [8]. Investigations include the measurement of electrical activity within cells, tissues, or organs, offering vital insights into the mechanisms governing physiological processes. By analysis of electrical signals from experiments, the complex interplay of ion channels and membrane potentials can be understood, providing crucial information on cellular function and communication. For example, the work of Hodgkin and Huxley showed how the membrane potentials of giant squid are related to their ion channels and the models developed have been widely applied to neuronal, sensory and cardiac cells [17]. Electrophysiology experiments serve as an indispensable tool in biomedical research and contribute to the development of innovative treatments for a wide range of neurological and cardiovascular conditions. It is hoped this will also be possible with bacterial infections [8] [18] [19] [20] [21].

Since bacteria are excitable matter, both individually and as biofilms, we hypothesised that there could be neuronal-like responses to electrical stimulation and such behaviour was previously claimed by Stratford et al [22] for planktonic bacteria. In general, the applied membrane potential due to AC stimulation of spherical cells by external electrodes can be predicted from the Schwann equation [23] [24], where the transmembrane voltage Δ*ψ*_*m*_ is

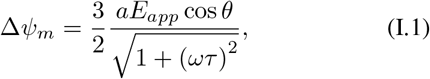

where *a* is the radius of a cell (assumed spherical), *E*_*app*_ is the applied electric field strength, *θ* is the angle between the electric field and the normal from the centre of the sphere to a given point on the cell membrane, *ω* is the angular frequency of the AC field, i.e. *ω* = 2*πf*, and *f* is the frequency of the field. *τ* is a time constant given by

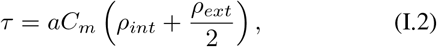

where *C*_*m*_ is the membrane capacitance, *ρ*_*int*_ is the internal fluid resistivity and *ρ*_*ext*_ is the external fluid resistivity.

Our article has two key contributions. First, the application of an AC electric field was not observed to give rise to spiking potentials, in direct contradiction to the study of Stratford et al in PNAS [22]. Instead, the dips in the intensity of cationic fluorophores on application of the AC electric field could be adequately explained in terms of AC electro-osmosis, a microfluidic phenomenon well established in the microfluidic literature [25] [26] and independent of bacterial electrophysiology. Second, we explored transport of a range of molecules and nanoparticles through bacterial biofilms driven by AC electro-osmosis. This has not been previously observed in the literature and provides a novel mechanism to increase the transport of particles through the biofilm structure. It could be used to treat persistent biofilm infections in tandem with antibiotics to increase their efficacy.

### A. AC Electro-Osmosis

AC electro-osmosis involves the microfluidic response of charged particles in a fluid to an applied AC electric field [27] [28]. Diffuse double layers form on the surface of conducting electrodes and their equilibriation time can be comparable with that of AC electric field oscillations [25]. Microfluidic pumping due to electro-osmosis occurs if the frequency of the applied AC field corresponds to the time for the double layers to form on the electrodes. When there are two co-planar electrodes, an AC current will generate a fluid flow which is dependent both on the dimensionless frequency (Ω) and on the applied voltage (*V*). The slip velocity (*v*_*s*_) of these particles is

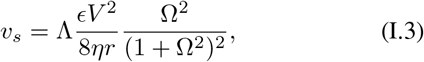

where *η* is the viscosity of the fluid, *ϵ* is the permittivity of the medium, *r* is the particle position on the electrode surface and Λ is

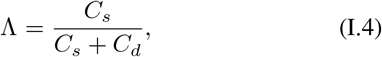

where *C*_*s*_ is the Stern capacitance per unit area and *C*_*d*_ is the diffuse layer capacitance per unit area. Ω can also be expressed by

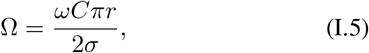

where *ω* is the applied frequency, *C* is the capacitance (given by Λ*E/λ*_*D*_, where *λ*_*D*_ is the Debye length), and *σ* is the conductivity.

In the current investigation, AC electro-osmosis masked any phenomena involving the electrophysiology of the bacteria by transporting fluorescent dyes towards the electrodes which were supplying the electric field. The dyes were then partially hidden from view on the electrodes. AC electro-osmosis therefore modulated the overall brightness of the region of interest in the microscope via two mechanisms: by concentrating the fluorophores in certain regions, e.g. the stagnation points of the microfluidic flows, and driving adsorption of the fluorescent dyes on to the electrodes.

## II. RESULTS

### A. Single Cells

We were motivated by our previous experiments that showed that blue light triggers electrical hyperpolarization in *E. coli* cells, *B. subtilis* cells and their biofilms [7] [14]. Knowing the cells and biofilms respond to blue light stress, the electrophysiological responses which involve electrical stress could be studied. Inspired by Stratford et al [22], bespoke microfluidic apparatus were created with two gold electrodes separated by a small gap (from 100 to 500 µm), which contained a liquid inoculum of *E. coli* labelled with Thioflavin-T (ThT) in a polydimethylsiloxane (PDMS) microfluidic device (Figure II.1). A burst of AC voltage was then applied across the electrodes of 10 V peak-to-peak (pp), to compensate for the fact that the gap was slightly wider than Stratford et al [22] to provide similar electric fields (see Methods in section IV).

**FIG. II.1.**
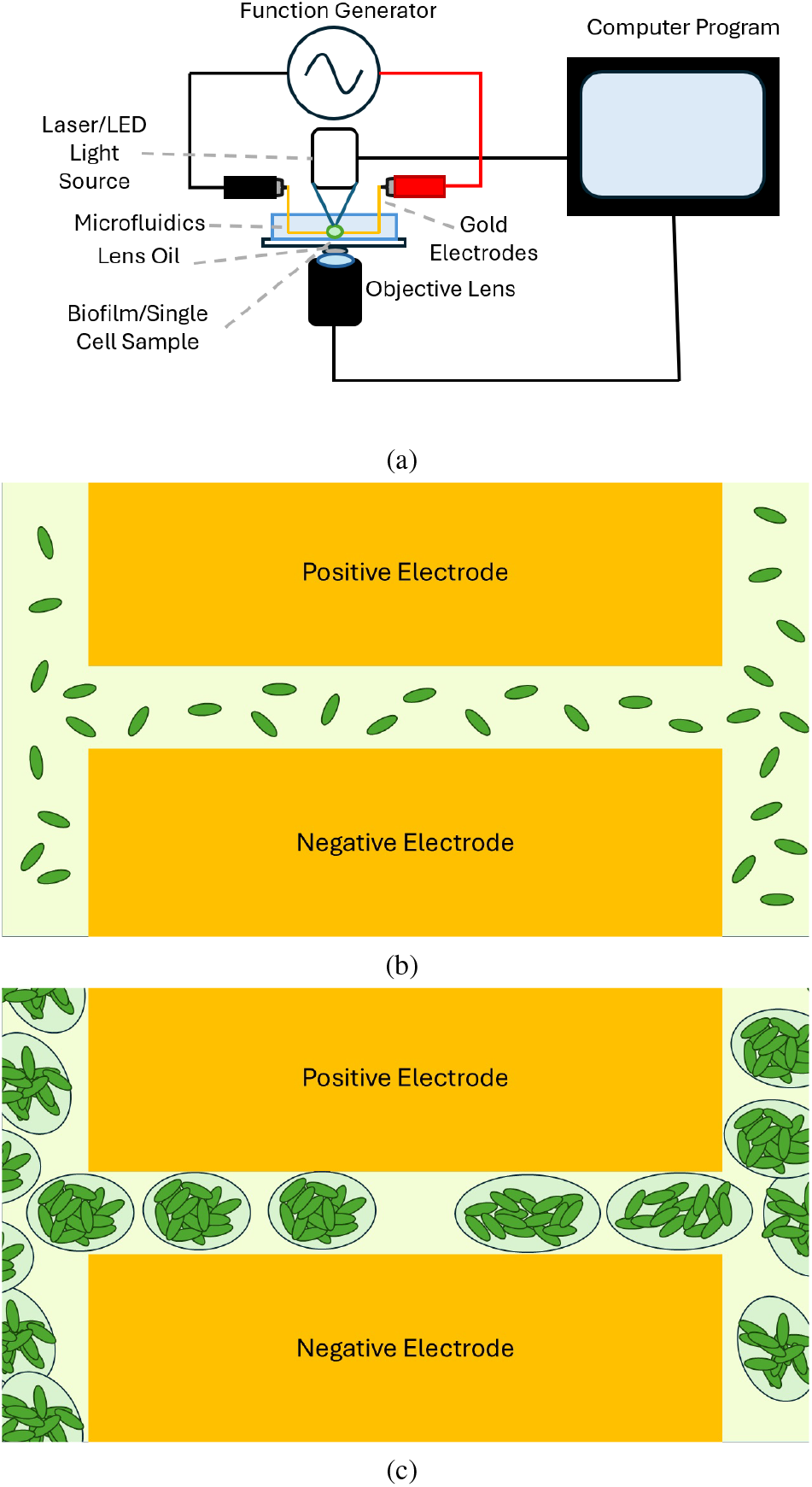
(a) Schematic diagram of the setup used in the experiment. Crocodile clips connect the function generator to two gold electrodes that have a gap between them where the sample resides. The microscope and light source are controlled via a computer program. (b) and (c) are schematics of bacteria residing between the two electrodes in the planktonic state and the biofilm state respectively viewed from the top. The bacteria are shown as a green ellipsoids and the electrodes are shown as two yellow rectangles. The background of the image is coloured to represent the media. In (c) the biofilms are surrounded by an extracellular matrix represented by an ellipsoid. The distance between the gap and the size of the bacteria are not to scale. The gap is around 200 µm and bacteria should be around 1 µm.

There is a characteristic decrease in fluorescence when the AC electric field is applied to a suspension of *E. coli* with ThT placed between the electrodes (Figure II.2).

The experiments were repeated using another fluorophore, propidium iodiode (PI), and a mixture of PI and ThT. The bacteria were also stimulated with more electric pulses to see if there are additional electrophysiological phenomena, such as resonance, which is present in the spiking dynamics of some neurons [29] [30].

**FIG. II.2.**
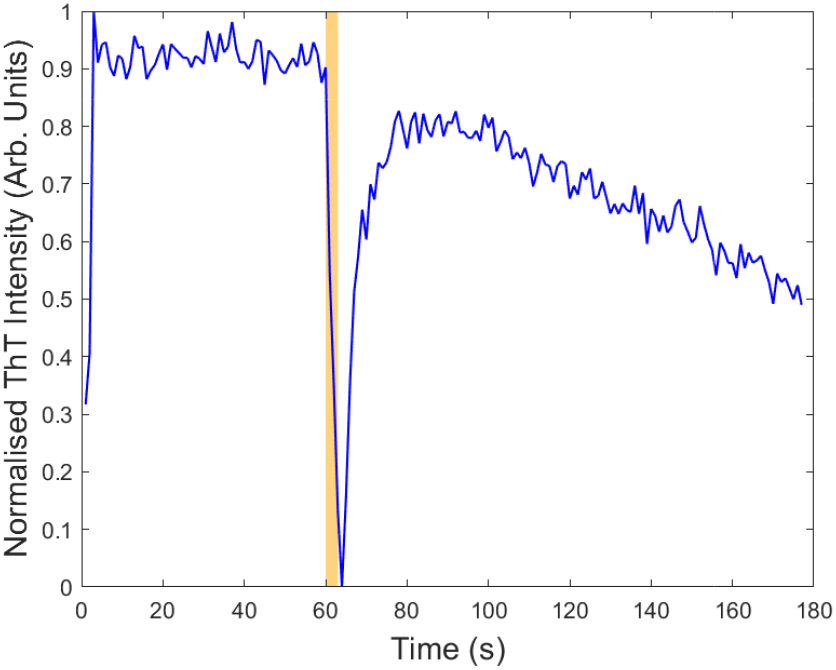
Fluorescence intensity (shown in blue) as a function of time for one AC electric field pulse applied to an *E. coli* suspension stained with ThT after one minute of no electrical stimulus. The pulse duration was 3 seconds and is highlighted in orange in the graph. The voltage was 10 V pp and the AC frequency was 1 kHz. Measurements were taken in an Olympus IX83 inverted microscope (Klaus Decon Vision) using blue Lumencor LED excitation.

PI alone was used to label the *E. coli* suspensions, Figure II.3 (a), and the dips in fluorescence were still present. Furthermore, when labelling with both ThT and PI, Figure II.3 (b), there is a dip in fluorescence in both fluorescence channels. The results were consistent for both PI and ThT. If electroporation was occurring [31], the PI would enter the cells and increase their brightness. Instead, the same characteristic dips were observed for both PI and ThT.

**FIG. II.3.**
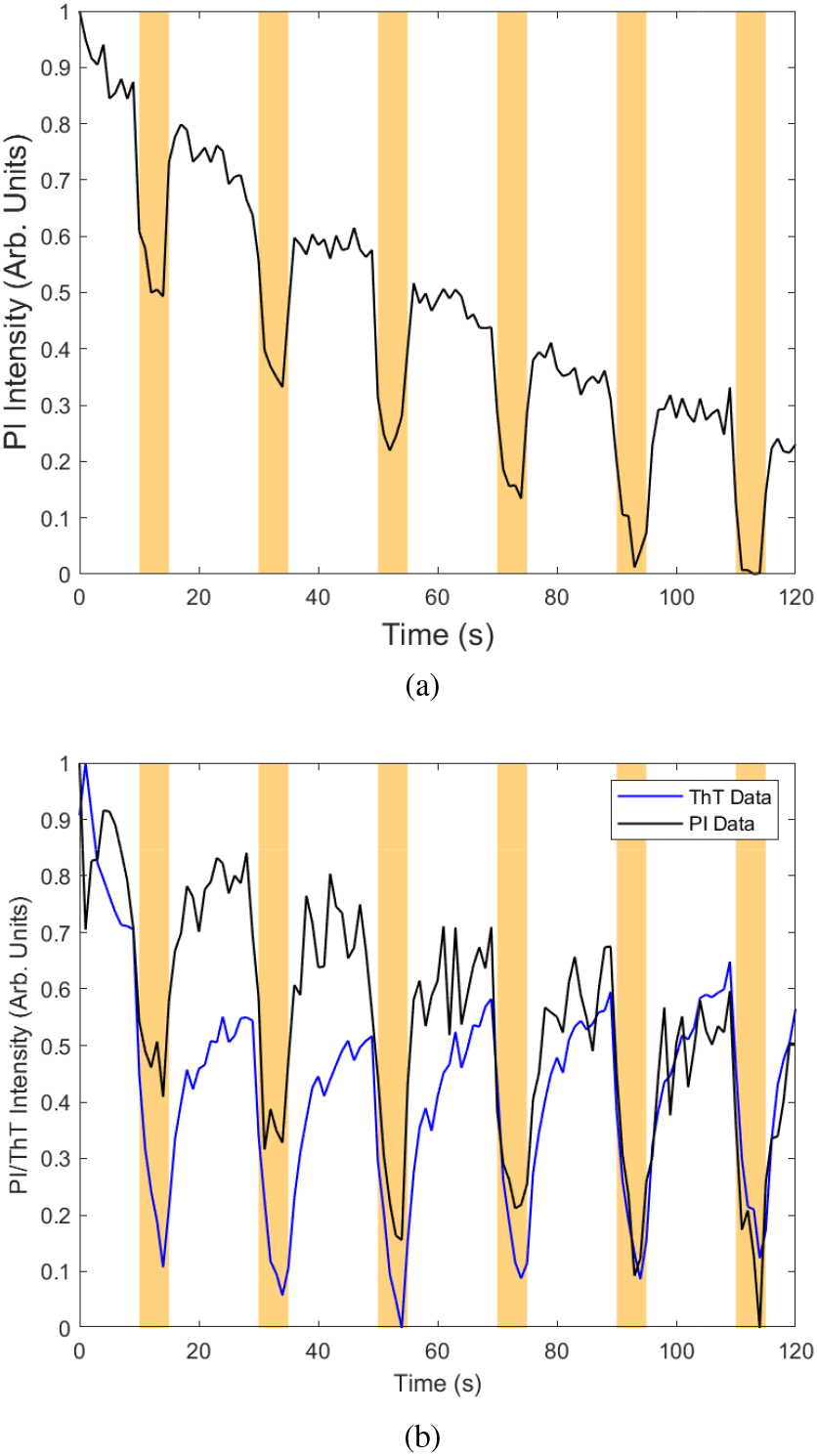
Fluorescence data of 6 AC electric pulses applied to an *E. coli* suspension stained with PI after 10, 30, 50, 70, 90 and 110 seconds. The duration of the pulses were 5 seconds and this period is highlighted in orange in the graph. The voltage was 10 V pp and frequency was 1 kHz. (a) Shows when only PI was present in black and (b) shows when both PI and ThT were present in the sample. Measurements were taken in an Olympus IX83 inverted microscope (Klaus Decon Vision) using blue Lumencor LED excitation.

The average fluorescence was calculated by integrating all of the region of interests in the fluorescence images. However, the same behaviour is also observed if only the cells were highlighted to analyse the brightness (Figure II.4 and Figure II.5 for ThT and PI respectively).

**FIG. II.4.**
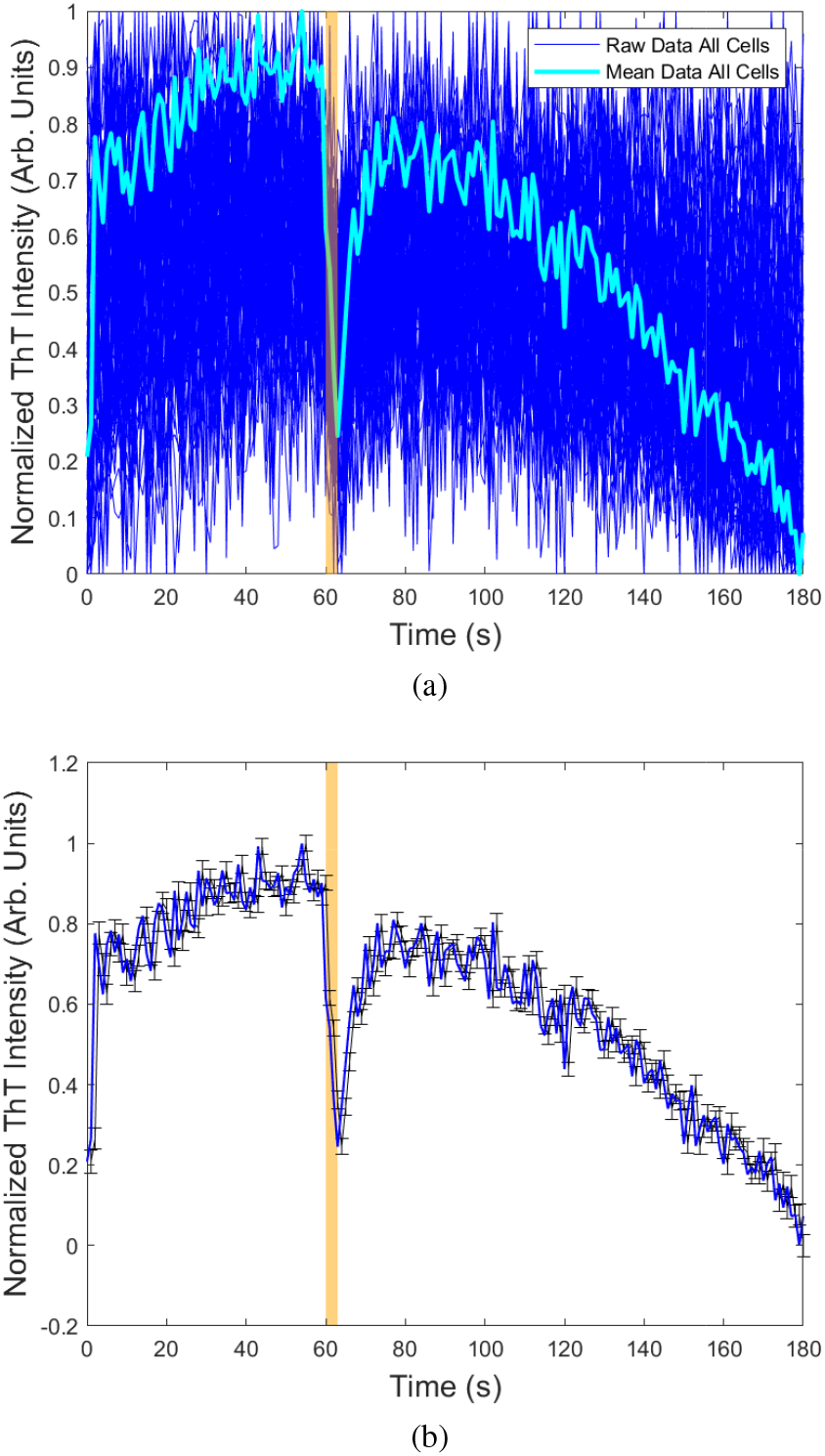
Fluorescence data of 6 AC electric pulses applied to the *E. coli* suspensions stained with ThT after 10, 30, 50, 70, 90 and 110 seconds. The duration of the pulses were 5 seconds and this period is highlighted in orange in the graph. The voltage was 10 V pp and frequency was 1 kHz. (a) All of the data for all single cells in blue and the mean of these points in light blue. (b) Mean data for single cells in blue with the corresponding error bars for each plot point. The error bars correspond to the standard error. Measurements were taken in an Olympus IX83 inverted microscope (Klaus Decon Vision) using blue Lumencor LED excitation.

**FIG. II.5.**
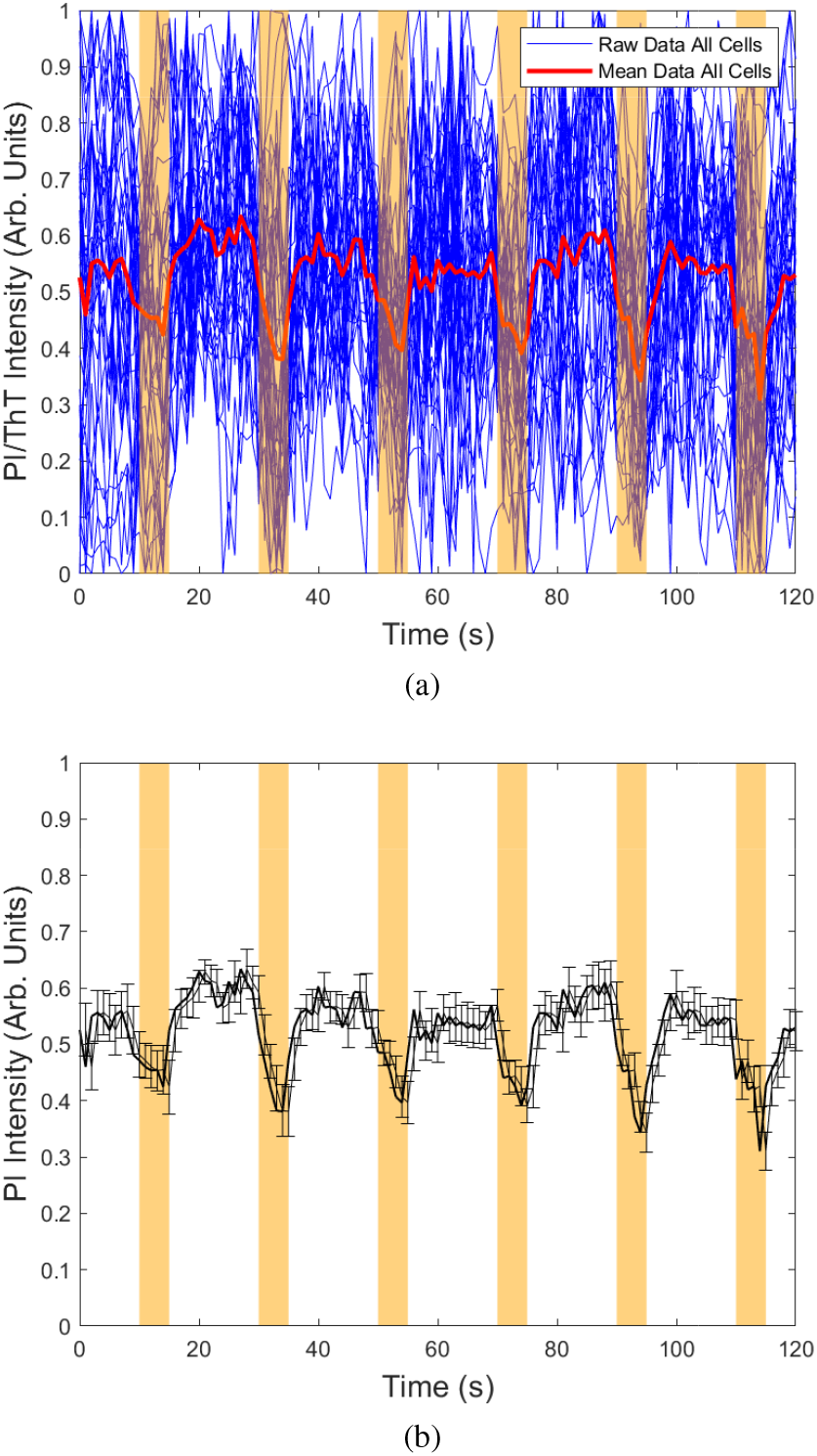
Fluorescence data of 6 AC electric pulses applied to the *E. coli* suspension stained with PI after 10, 30, 50, 70, 90 and 110 seconds. The duration of the pulses were 5 seconds and this period is highlighted in orange. The voltage was 10 V pp and frequency was 1 kHz. (a) Shows the data for all single cells in blue and the mean of these points is in red. (b) Mean data for single cells in blue with the corresponding error bars for each plot point. The error bars correspond to the standard error. Measurements were taken in an Olympus IX83 inverted microscope (Klaus Decon Vision) using blue Lumencor LED excitation.

A combination of PI and Syto9 is commonly used for live-dead assays [32]. Interestingly, Figure II.6 (a), shows the same dips in fluorescence are observed with Syto9. Finally, a fourth fluorophore, Sytox Green, was used due to its ability to permeate dead cells or membrane compromised cells better than PI [33]. Similar to the previous experiments with PI, the fluorescence also decreased when an electric pulse was applied in the presence of Sytox Green, Figure II.6 (b).

**FIG. II.6.**
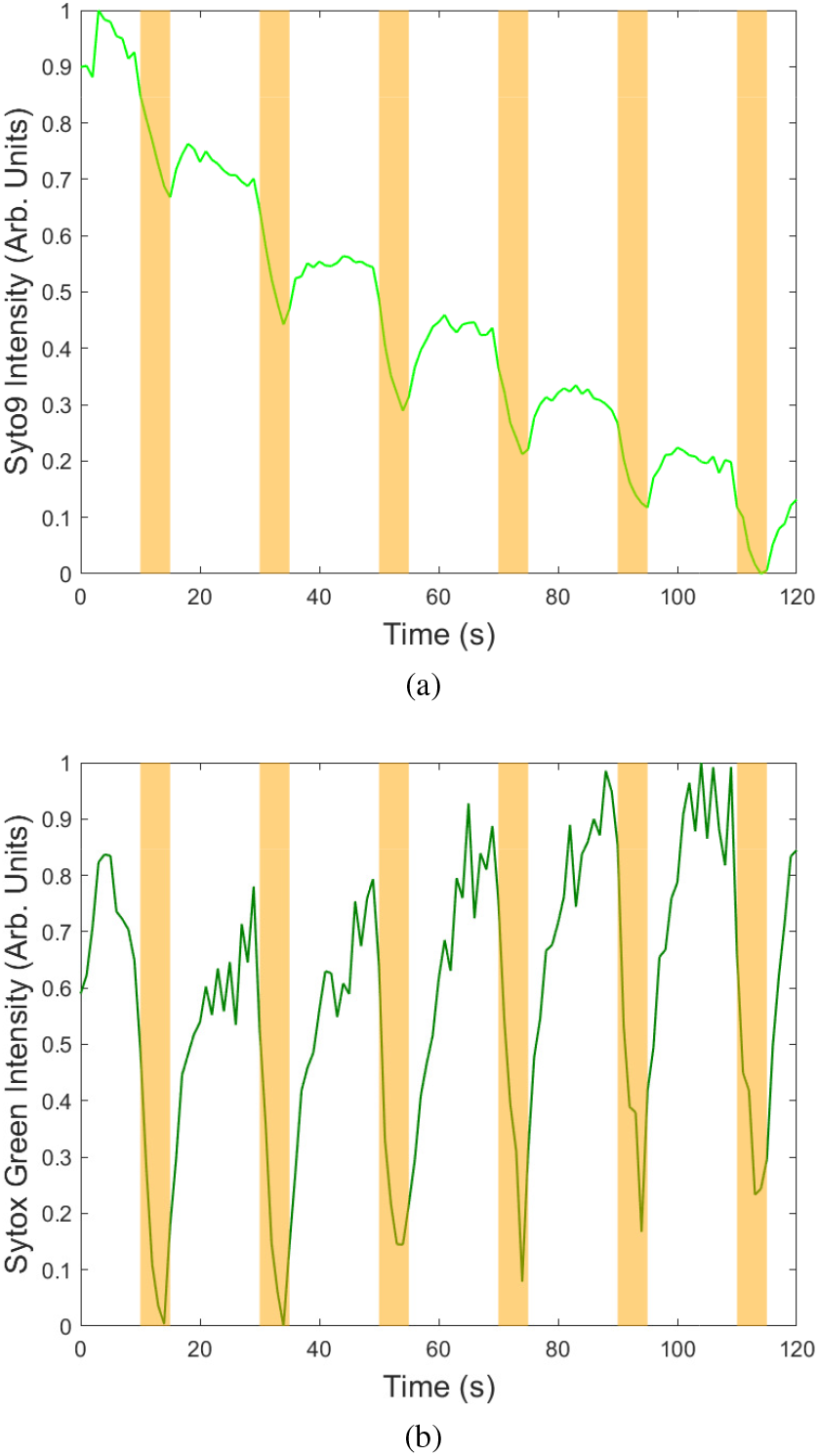
Fluorescence data of 6 AC electric pulses applied to the *E. coli* suspension stained with ThT after 10, 30, 50, 70, 90 and 110 seconds. The duration of the pulses were 5 seconds and this period is highlighted in orange in the graph. The voltage was 10 V pp and the frequency was 1 kHz. (a) Fluorescence data for Syto9 (green). (b) Fluorescence data for Sytox Green (dark green). Measurements were taken in an Olympus IX83 inverted microscope (Klaus Decon Vision) using blue Lumencor LED excitation.

The growth media for the *E. coli* was then changed from Lysogeny Broth (LB) to Davis Minimal Media (DM) with the addition of Casamino acids (CAA) to see whether the electrolyte is important. In Figure II.7 the fluorescence dips are still prominent when using ThT in DM.

**FIG. II.7.**
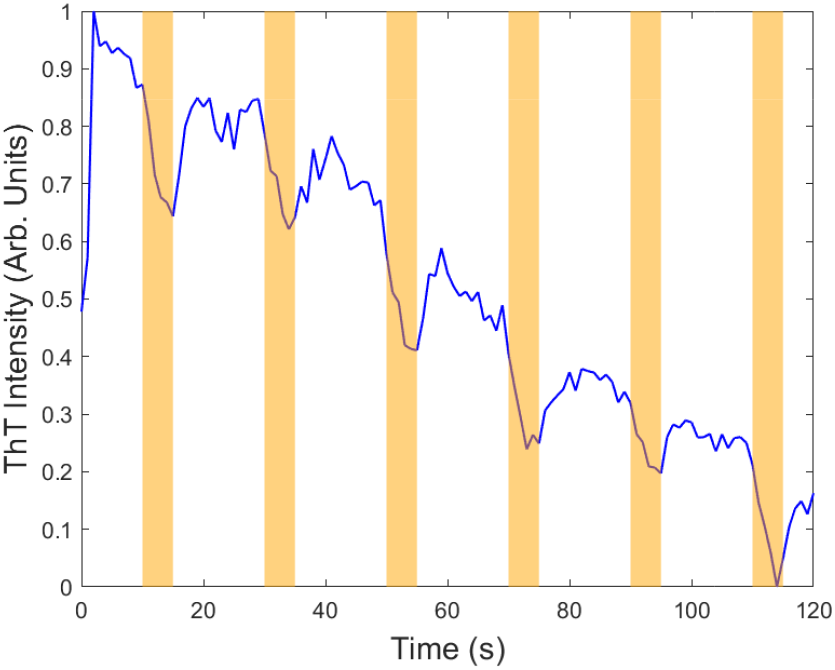
Fluorescence data (blue) of 6 AC electric pulses applied to the *E. coli* suspension stained with ThT with DM and CAA media after 10, 30, 50, 70, 90 and 110 seconds. The duration of the pulses was 5 seconds and this period is highlighted in orange in the graph. The voltage was 10 V pp and frequency was 1 kHz. Measurements were taken in an Olympus IX83 inverted microscope (Klaus Decon Vision) using blue Lumencor LED excitation.

A final check to see if the cells were responsible for the phenomenon of decreased fluorescence, was to repeat the experimental procedures with ThT and Sytox Green without any bacterial cells. Surprisingly, the fluorescence dips were still present without any cells and with the different media, Figure II.8 (a), Figure II.8 (b) and Figure II.8 (c).

To summarise, Figure II.9 shows the characteristic dips in fluorescence are independent of the fluorophore and independent of the presence of the cells i.e. it is not a biological phenomenon.

**FIG. II.8.**
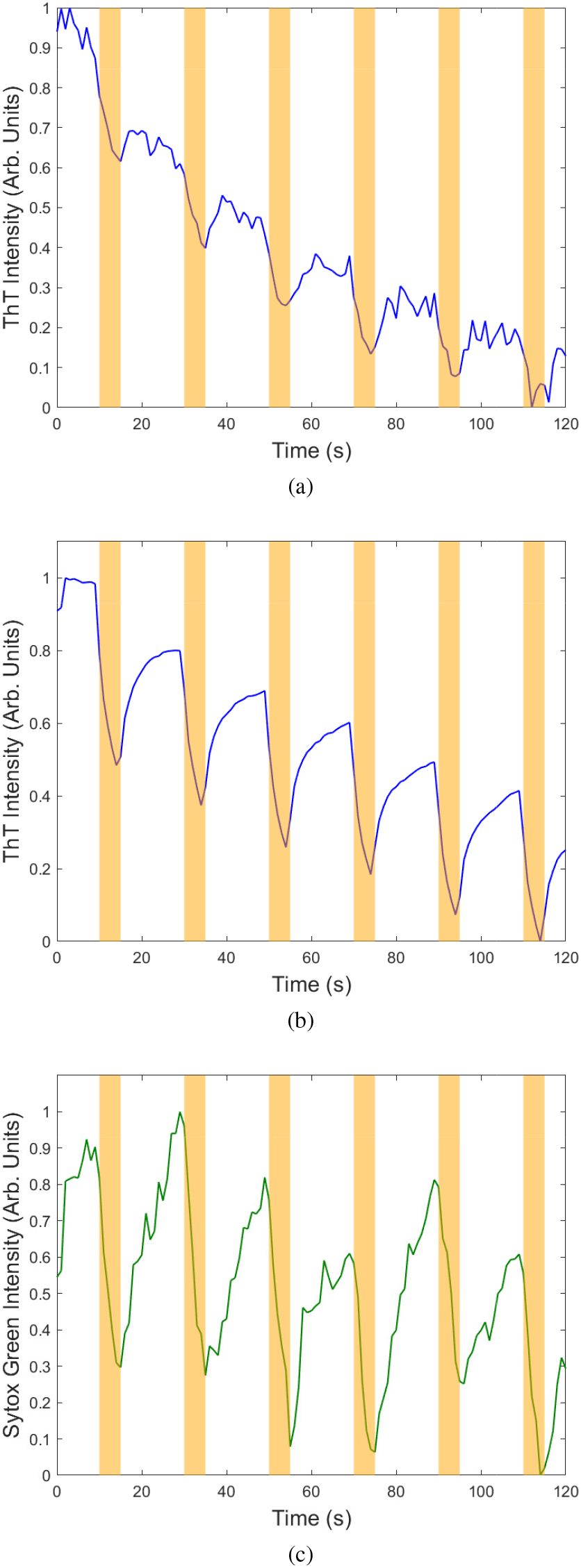
Fluorescence data of 6 AC electric pulses applied to the media without *E. coli* cells with ThT after 10, 30, 50, 70, 90 and 110 seconds. The duration of the pulses were 5 seconds and this period is highlighted in orange. The voltage was 10 V pp and frequency was 1 kHz. (a) Shows fluorescence data in blue for ThT without cells in LB. (b) Shows fluorescence data in blue for ThT without cells in DM and (c) shows fluorescence data in dark green for Sytox Green without cells in DM. Measurements were taken in an Olympus IX83 inverted microscope (Klaus Decon Vision) using blue Lumencor LED excitation.

**FIG. II.9.**
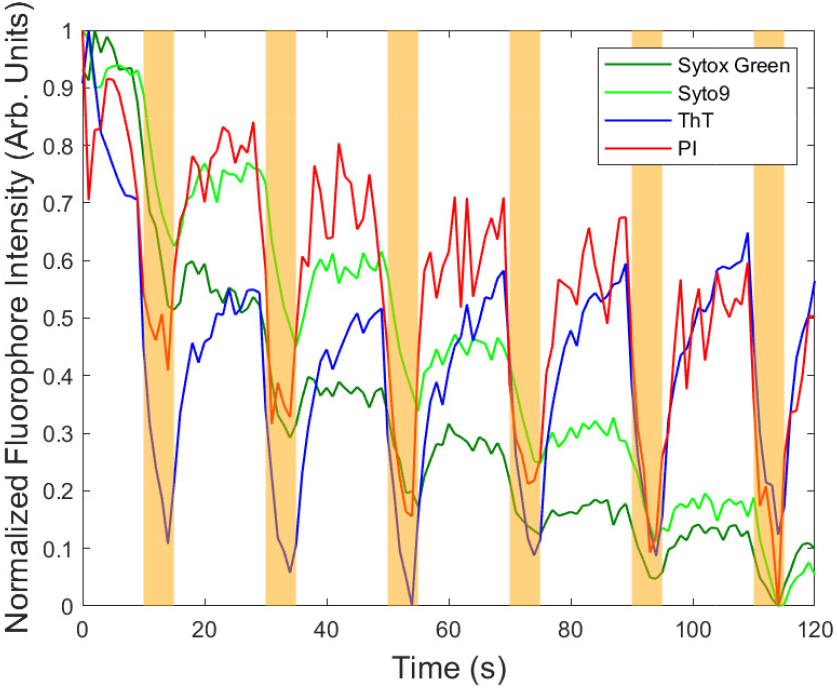
Fluorescence data of 6 AC electric pulses applied to the media containing each type of fluorophore without *E. coli* cells after electrical stimulation at 10, 30, 50, 70, 90 and 110 seconds. The duration of the AC pulses was 5 seconds and they are highlighted in orange. The voltage was 10 V pp and frequency was 1 kHz. Measurements were taken in an Olympus IX83 inverted microscope (Klaus Decon Vision) using blue Lumencor LED excitation.

The Schwann equation (equation I.1) predicts an electrical response of the membrane in the cells under AC stimulation. However, a slip velocity is also expected created by AC electro-osmosis, equation I.3. The decrease in fluorescence is proportional to *V*^2^, as predicted by equation I.3 (see supplementary information Figures II.1 to II.5, which are in reasonable agreement with the predicted scaling) and we thus conclude that the dips in the fluorescence (phantom action potentials) are solely due to AC electro-osmosis, masking any effects due to electrical stimulation of the bacteria.

It is possible to simulate AC electro-osmosis from the two linear electrodes and we did so for our geometry as shown in Figure II.10 and the Stratford et al [22] geometry (see Supplementary Information). The simulation involved solving the Stokes equation coupled with solving Laplace’s equation for the potential using the finite element method alongside boundary conditions with the Helmoltz-Smoluchowski formula (See Supplementary Information for full description of the equations and methods used). The simulations are based off of the work of Green et al, which showed that the AC electro-osmosis effects are numerically calculable and agree with experimental data [34]. The slip velocity is then calculated alongside the electric field components. The fluids are driven by the electro-osmosis flows and fluorophores will accumulate in different regions modulating their concentrations and causing the dips.

**FIG. II.10.**
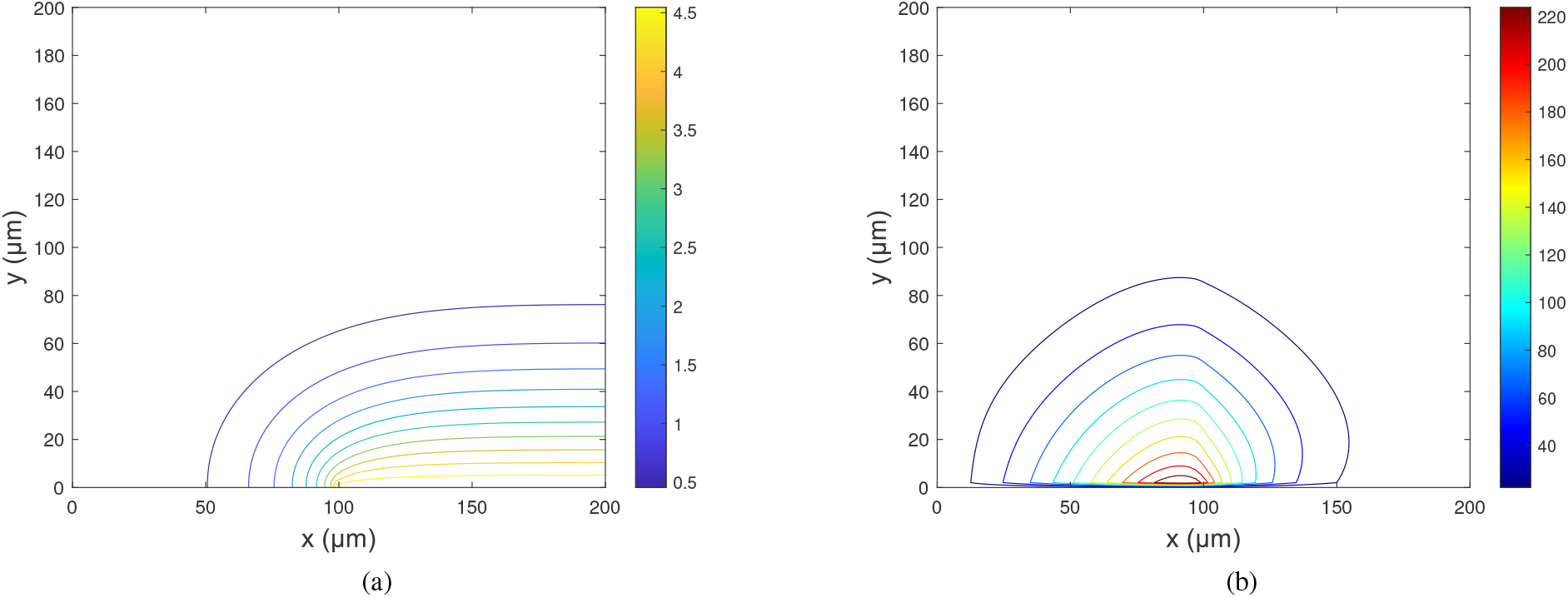
Microfluidic simulation of AC electro-osmosis from two linear electrodes separated by a gap of 0.2 mm with a voltage of 10 V and a frequency of 1000 Hz. The view is from the side of one of the two electrodes (which have identical behaviour) similar to the electrode setup shown in Figure II.1 (a). A finite element model is solved based on a combination of Stokes equation and the Laplace equation alongside the Helmholtz-Smoluchowski formula (Supplementary Information). (a) Shows the electrical potential with the colour-bar in Volts and (b) shows the streamline flow contour plot for the fluid mechanics. The colour-bar is in µm/s. There is a plane of mirror symmetry at x = 0 for both (a) and (b)

### B. Biofilms

Biofilms were grown with ThT (as originally used in our light stimulation experiments [7] [14]).The fluorescence dips were still prominent with bacterial biofilms (Figure II.11). AC electro-osmosis drives the flurophores through the EPS of the biofilms and around the bacterial cells.

**FIG. II.11.**
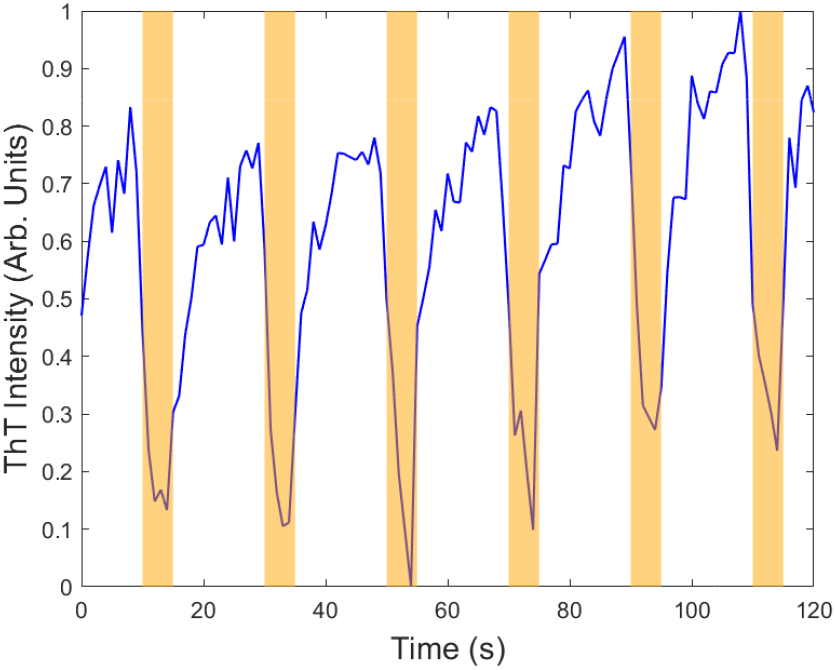
Fluorescence data (blue) of 6 AC electric pulses applied to *E. coli* biofilm in LB stained with ThT after 10, 30, 50, 70, 90 and 110 seconds. The duration of the AC pulses 5 seconds and this period is highlighted in orange. The voltage was 10 V pp and frequency was 1 kHz. Measurements were taken in an Olympus IX83 inverted microscope (Klaus Decon Vision) using blue Lumencor LED excitation.

#### 1. Fluorescent Beads

Motivated by the experiments on the small molecule fluorophores, we questioned whether bigger particles could also be driven through biofilms. Sulfate modified polystyrene colloidal spheres of average size 100 nm and carboxylate modified polystyrene colloidal spheres of average size 30 nm were chosen as they have negative charges. ThT was used alongside the beads for visualisation of the bacteria. In general, the EPS of biofilms also have negative charges, so large scale electrostatic complexation is not expected and the beads could be free to move through the biofilms.

The electrical stimulation experiment was repeated, with the spheres added to the ThT (see Methods section). The 30 nm carboxylate beads show the characteristic dip in fluorescence intensity when the electric pulse is applied similar to the ThT (Figure II.12). The beads’ fluorescence intensity appears more noisy due to the relative fluorescence yields of the particles, but the average behaviour is clear.

**FIG. II.12.**
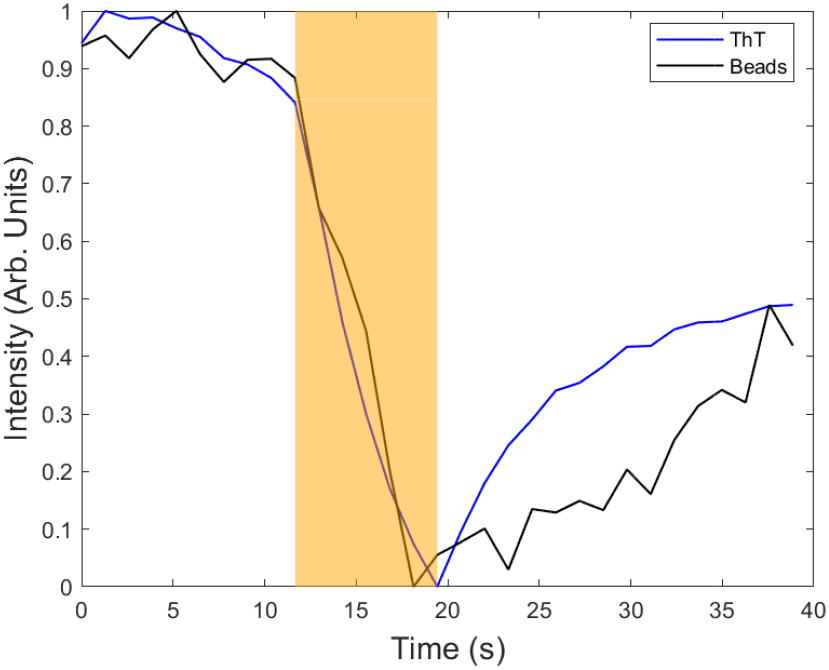
Fluorescence data shown in blue and black of ThT and beads respectively, for 30 nm carboxylate coated beads with an AC electric pulse after 10 seconds in an *E. coli* biofilm section. The pulse was 5 frames long with each frame being 0.772 seconds.The voltage was 10 V pp and frequency was 1 kHz. Measurements were taken in a Leica TCS SP8 AOBS inverted gSTED confocal microscope.

The 100 nm sulfate coated beads also show the characteristic dip in intensity when the electric pulse is applied (Figure II.13). However, different lag times were observed in the two fluorescence channels for the ThT and beads for some regions of interest, Figure II.13 (b). This suggests that the ThT and the beads are independent of one another during transport through the biofilm. The short sedimentation time of 1 *µ*m polystyrene beads meant there were practical challenges to perform the control experiment without biofilm, since no beads were observed in the bulk fluid. The 1 *µ*m beads did not move when added to the biofilm, presumably because they were jammed in to the polymeric network of the EPS.

**FIG. II.13.**
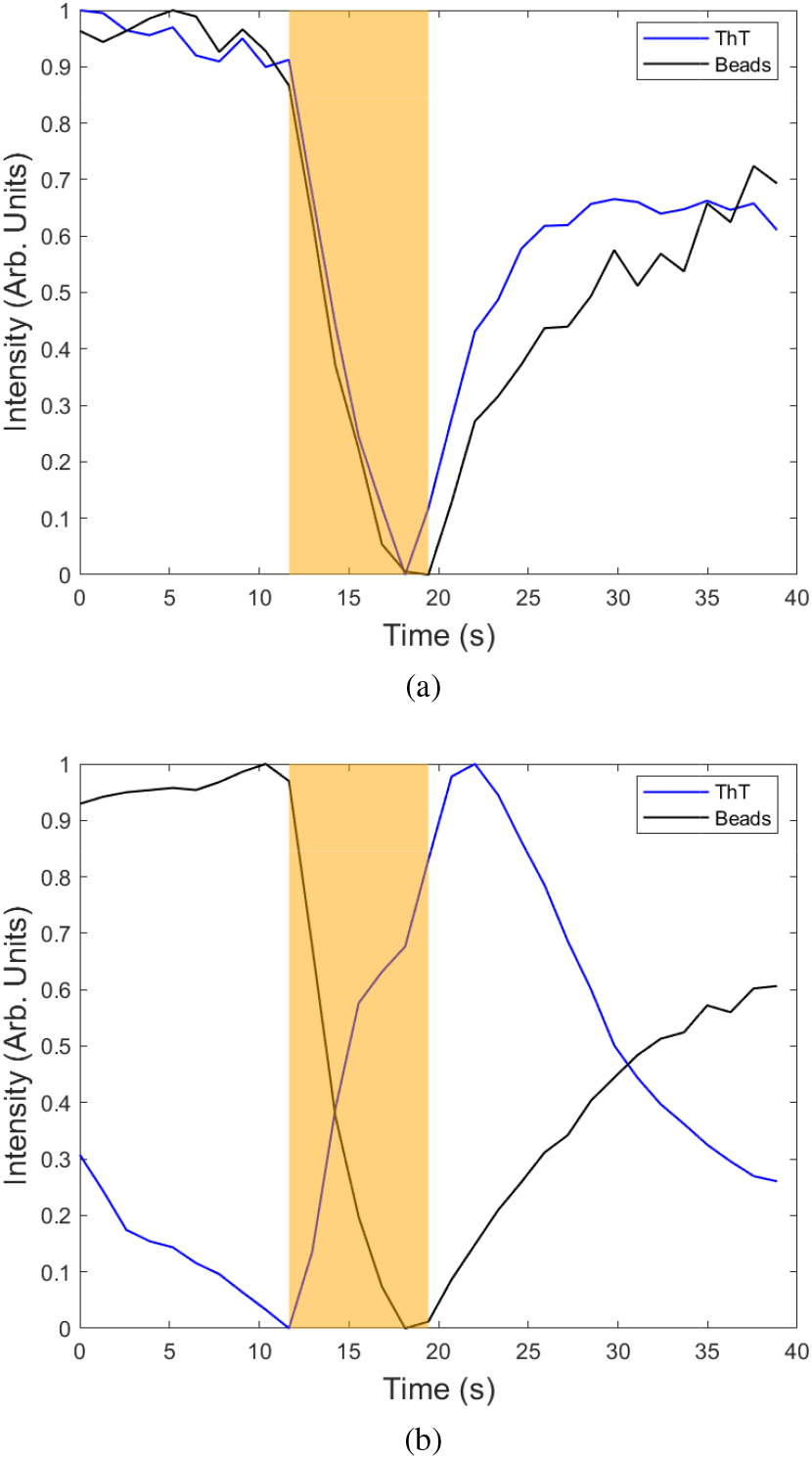
Fluorescence data of an AC electric pulse applied to an *E. coli* biofilm section with ThT and 100 nm sulfate coated beads after 10, seconds for a result of 5 frames. Each frame was 0.772 seconds long. The voltage was 10 V pp and frequency was 1 kHz. (a) Shows fluorescence data for ThT (blue) and the beads (black) in a specific region of interest. (b) Shows fluorescence data for ThT (blue) and the beads (black) for a different region of interest. Measurements were taken in a Leica TCS SP8 AOBS inverted gSTED confocal microscope.

#### 2. DNA

The transport of DNA through biofilms is biologically interesting because extracellular DNA (eDNA) is a charged molecule with structural role in biofilms [35], naturally competent bacteria can uptake eDNA as short as 20bp [36], and DNA can be used as a nutrient [37]. PCR was used to produce fluorescent monodisperse DNA labelled with the fluorophore Cy3. Care was taken to remove the unbound fluorophores and thus any background signal.

500 base pair (bp) DNA was used. DNA molecules are known to experience AC electro-osmosis in free solutions which has been used to modulate their concentration [38].

The end-to-end size of the DNA in solution can be calculated from the worm-like chain model. The end-to-end length (*R*) is

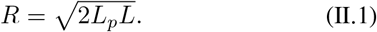

where *L*_*p*_ is the persistence length of DNA (under standard conditions *L*_*p*_ ≈ 50 nm) and *L* is the contour length of the DNA. *L* can be approximated by the number of base pairs multiplied by their size, 0.34 nm. Therefore, for a 500 b.p. DNA, *R* is estimated to be 131 nm.

The DNA showed characteristic dips in fluorescence intensity, (Figure II.14) as it moved through the *E. coli* biofilms. The DNA also showed the characteristic dips in intensity with repeated electrical stimulation, Figure II.15. The two fluorescence channels suggest that the Cy3 labelled DNA was independent of the ThT as there are clear differences in the intensity for each normalised curve.

**FIG. II.14.**
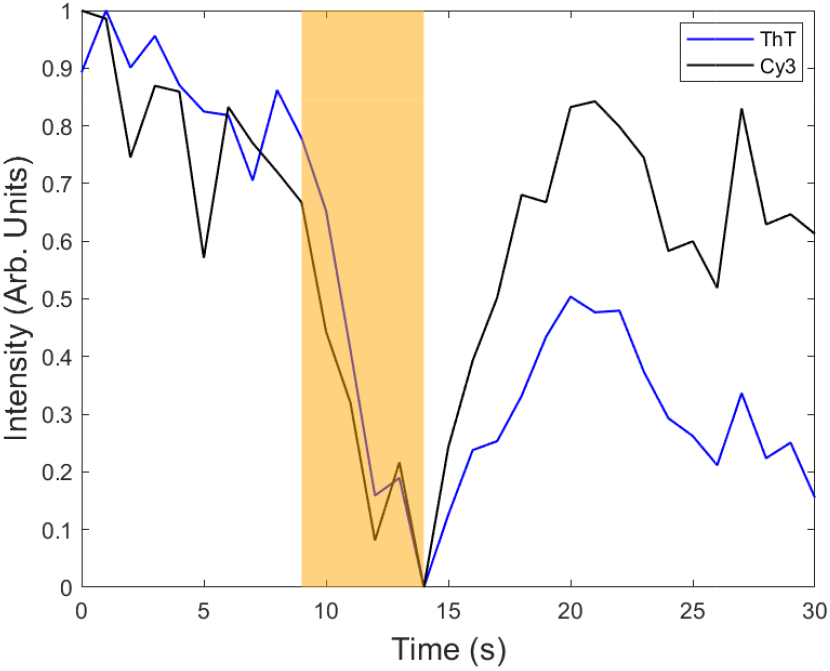
Fluorescence data shown in blue and black of ThT and Cy3 respectively, for 500 bp DNA with an AC electric pulse after 10 seconds in an *E. coli* biofilm section. The AC electric pulse was 5 frames long with each frame being 1 second. The voltage was 10 V pp and frequency was 1 kHz. Measurements were taken in a Leica TCS SP8 AOBS inverted gSTED confocal microscope.

**FIG. II.15.**
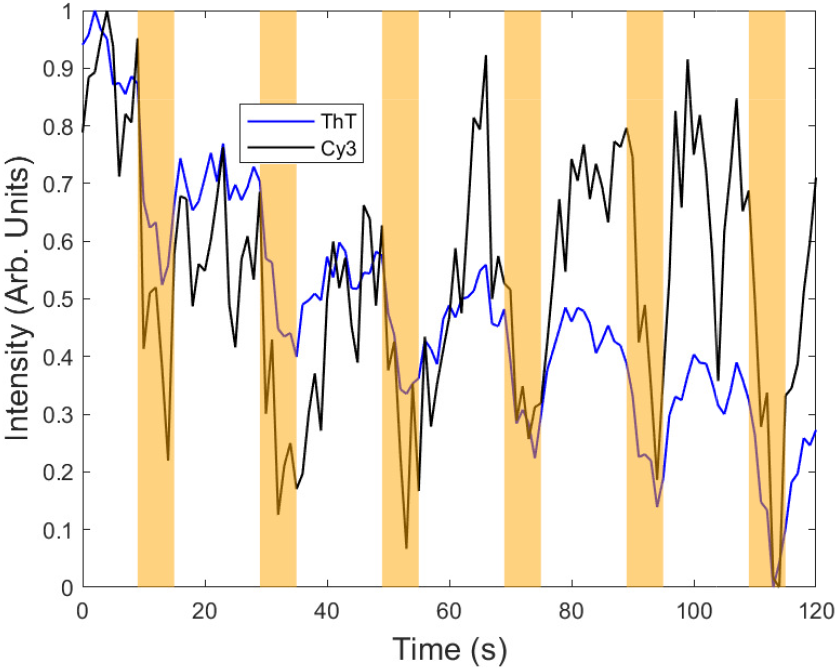
Fluorescence data shown in blue and black of ThT and Cy3 respectively, for 500 bp DNA with an AC electric pulse after 10, 30, 50, 70, 90 and 110 seconds in an *E. coli* biofilm section. Each AC electric pulse was 5 frames long and each frame was 1 second. The voltage was 10 V pp and the frequency was 1 kHz. Measurements were taken in a Leica TCS SP8 AOBS inverted gSTED confocal microscope.

### C. Time Constant Measurements

To quantify AC electro-osmosis for different fluorophores, an exponential function was fitted to the dips in the fluorescence. The fluorescence intensity (*F*) as a function of time was described by the empirical equation,

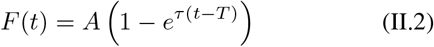

where *A* is the normalised amplitude of the fluorescence decay, *τ* is the relaxation rate, *t* is the time and *T* is the initial time for the dip to start recovering fluorescence. Fits were obtained for the fluorescent dips in the different conditions and the average of the variables were taken.

## III. DISCUSSION

Although the gap between the electrodes varied in length slightly from experiment to experiment (varying the electric field between 100 and 500 V/cm), the same phenomenon was observed in all trials. There was a prominent dip in fluorescence whenever an electric pulse was applied. This differed from Stratford et al’s results [22]. Furthermore, Stratford and co-workers claimed there was no effect of light on hyperpolarization of cells in contradiction to our previous results [7]. Stratford et al attribute the dips in fluorescence to hyperpolarization of the cells in response to electrical stimulation. In contrast, we find the fluorescent dips are satisfactorily explained by AC electro-osmosis; a microfluidic effect [25] [26]. The PDMS in the microfluidic chamber does contribute to the low fluorescent background present in the videos, but it was uniform throughout the regions of interest and did not affect the overall observed results.

An initial hypothesis was that the decrease in fluorescence when an electrical pulse was applied, followed by the subsequent increase in fluorescence, was due to reversible electroporation of the bacterial cells. Electroporation can occur in *E. coli* when electric fields are applied in the range 0.5 to 4 kV/cm and it is a standard technique to transform DNA chains into bacterial cells [31]. The electric field strength used in the experiments was on the order of 10^2^ V/cm; slightly lower than the accepted electroporation thresholds (although it could depend on the exact membrane biochemistry of a particular bacterium). The magnitude of the electric field applied is predicted by the Schwann equation [23]. Reversible electroporation occurs when a threshold value of an electric field is reached. It causes small nanoholes to form in the membrane, compromising the integrity of the membranes and allowing molecules to travel in and out before the membrane heals via self-assembly [39] [31]. Therefore, as PI is only permeable to compromised cells, it was initially hypothesised that the cells would increase in brightness when an electric pulse was applied in the presence of PI, resulting in the dye passing through the previously impermeable membrane and complexing with the DNA of the cell. This was not observed with PI in our experiments (Figure II.3a).

Sytox Green only binds to dead cells or cells with compromised membranes similar to PI. Sytox Green and PI were previously used to investigate electroporation in cells [40]. Sytox Green also did not enter the cells during electrical stimulation, Figure II.6 (b). The phenomenon is thus unlikely to be electroporation, since the fluorescence in the cells does not increase.

The media the *E. coli* were grown in, Lysogeny Broth (LB), is known to have background autofluorescence, so the media could have influenced the fluorescence dips [41]. Davis Minimal Media with the addition of Casamino Acids (CAA) was therefore used for comparison, as it possesses little autofluorescence due to the high purity components used due to presence of only essential elements (a phosphate buffer with citrate, nitrogen, magnesium and thiamin). No significant changes in results were seen using the new media, so the effects of media autofluorescence are discounted.

Combined with the fluorescence dips observed without cells, all the results strongly suggest it is a purely physical phenomenon decreasing the fluorescence of the cells e.g. Figure II.2. AC electro-osmosis is proposed to explain the decrease of fluorescence. This would also explain the slight movement of the cells whenever an electric pulse is applied, and the movement of the brightness gradient of the fluorophores when there were no cells, due to microfluidic pumping of the media (see supplementary videos).

The fluorescence dips were also present in bacterial biofilms. This suggests that AC electro-osmosis facilitates the movement of molecules through the EPS.

Other photochemical effects, such as the Stark effect and fluorescence quenching, were also considered to explain the phenomenon of the decrease in fluorescence, but these effects change the fluorescence by very little in organic systems, so the effects were deemed too small to produce the pronounced fluorescence dips [42] [43] [44] [45] [46].

The fluorescence dips are a universal phenomenon for all of the fluorophores whenever an AC electric pulse is applied. We also believe that the steady decrease in fluorescence superposed on the dips is caused by the fluorescent molecules becoming stuck to the electrodes.

In Figure II.16, there is a change in the relaxation rates depending on the fluorophores and protocol used. For both PI experiments, the values for the relaxation rate (*τ*) are larger than the other fluorophores. PI is a cationic fluorophore with the highest molecular weight (668.45 g/mol) of the four fluorophores used and it required the smallest amount of time to relax. PI has a high molecular charge of +2. Interestingly, ThT has the lowest mass of all four dyes (318.86 g/mol), but does not have the lowest *τ* value. This could be due to the net charge of ThT is +1. Syto9 then has a lower *τ* value than ThT despite having a larger molecular weight (400 g/mol) and being positively charged. Sytox Green possesses a slightly higher *τ* (with a molecular weight of 600 g/mol) with a charge of +2. Overall this suggests that the mass and charge of the fluorophores only have a small influence on their relaxation times during AC electro-osmosis.

**FIG. II.16.**
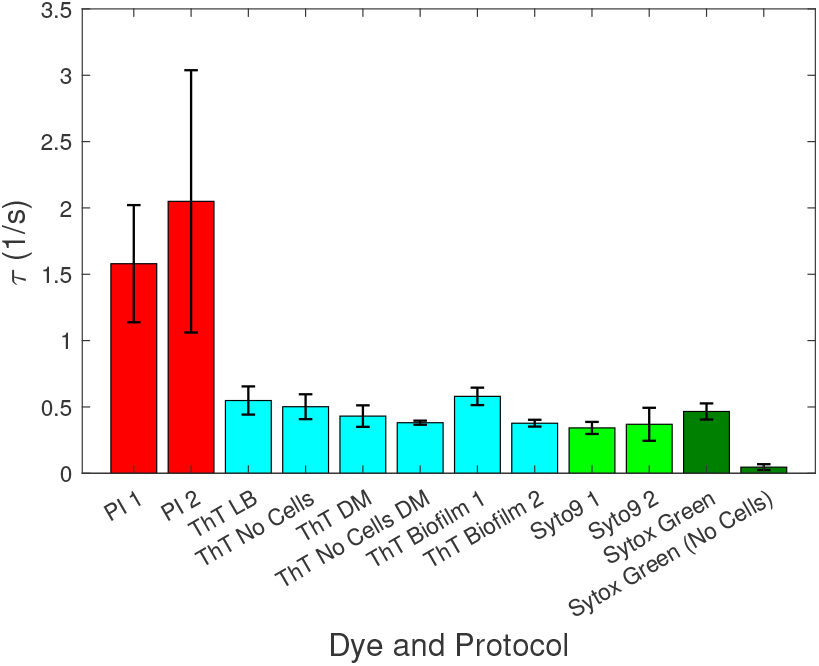
Relaxation rates (*τ*) of different fluorophores and protocols in *s*^*−*1^. The different fluorophores show different relaxation times. The error bars are the standard errors for multiple characteristic dips in fluorescence (*>* 10). PI is propidium iodide, ThT is Thioflavin T, LB is Lysogeny and DM is Davis Mignoli minimal media.

Furthermore, from the ThT data, there is no significant change of *τ* when there is a biofilm present compared with single cells or no cells. Nevertheless, there is a trend of decreasing relaxation rate for both ThT and Sytox Green when there are no cells present. The expression of biofilm does not significantly affect the electro-osmotic mobility of the fluorophores in agreement with previous studies on the diffusive mobility of fluorophores in biofilms [47].

An analogous artefact to that causing the phantom action potentials with bacteria may have affected the results presented by Cohen et al in 1969 using light scattering from neurons stimulated with electrodes [48]. The authors interpreted the dips in light scattering as due to changes in the neuronal membranes. An alternative explanation is that electro-osmosis of ions in the culture media caused electropumping and modulated the refractive indices of the solutions due to their microfluidics, which caused a change in the scattered light (proportional to the square of the refractive index difference). Thus artefacts due to microfluidic electropumping driven by AC or DC electro-osmosis may be relevant to a wide range of optical electrophysiology measurements due to their modulation of fluorophore or electrolyte concentration which will follow the driving electric fields [25]. Such phenomena will affect electrophysiology measurements during both DC and AC stimulation of excitable cells when they are investigated using fluorescence microscopy/spectroscopy, light scattering, birefringence and dichroism. Such optical methods are reasonably common in studies of the electrophysiology of stimulated neurons [49] [50] [51]. The use of intracellular recordings should reduce the magnitude of these artefacts and provides another advantage for optogenetics techniques. Furthermore, careful control experiments using identical microfluidic geometries and electrolyte concentrations may allow the effects of electro-osmosis to be separated from the electrophysiology of the cells.

Whether external electrical fields can stimulate bacterial cells to modulate their membrane potentials is still not well established. It is possible that the AC electro-osmosis is masking effects due to the bacteria in the fluorescence signals and they can be uncovered by a more judicious choice of microfluidic parameters e.g. electrode geometry and cell/electrode distances [52]. A possible solution would be to arrange the electrodes across a 2D slot or 1D capillary geometry to reduce the electro-osmotic flows while maintaining the stimulating electric fields across the bacterial cells. An alternative solution would be to investigate electrode coatings (e.g. spin coating thin layers of hydrophobic synthetic polymers such as PMMA), although this would inevitably reduce the electric field experienced by the bacteria and thus their response to electrical stimulation. A better solution would be to increase the viscosity of the growth media of the bacteria to reduce the AC electro-osmosis artefact (*v*_*s*_ ∼ *η*^−1^ in equation (I.3)) with the addition of viscosity modifiers e.g. glycerol or hydrophilic polymers, such as methylcellulose. The challenge in this case would be to ensure that the physiology of the bacteria is not significantly perturbed.

In general, electrode geometries could be optimized with future finite element modelling. Previous evidence for spiking potentials has been observed in a microelectrode array study on bacterial biofilms [53]. In the future it would also be useful to make detailed studies of electrical stimulation using optogenetic probes, since they can be confined within the bacterial cells reducing the effects of electro-osmosis [16].

The transport of molecules through biofilms needs more research, since it could be the rate limiting step of some antibiotics [5]. More extensive research has been performed on the transport of nanoparticles through hydrogels, such as mucus [54]. There is thus expected to be a charge and size filter on the motion of nanoparticles in biofilms in broad agreement with our experiments using AC electro-osmosis. However, more complex behaviour is possible in biofilms than mucin gels e.g. holes are created by *E. coli* in biofilms that will facilitate transport if the nanoparticles can permeate the lipopeptides that coat the holes [55].

## IV. METHODS

### A. Bacterial Strains

*E. coli DH5α* was used in all experiments. This strain is known to adhere well to surfaces and form strong biofilms

[56] [57].

### B. Microfluidics

The microfluidic device used to electrically stimulate the bacteria was built using polydimethylsiloxane (PDMS) with an elastomer kit (Conro Electronics Dowsil Sylgard). The PDMS was mixed with an elastomer curing agent with a 20:3 ratio and degassed for *>*1 hour. This ensures that there are no bubbles in the PDMS.

A 3D-printed template (see supplementary section) that is 35 mm wide, 60 mm long and 5 mm tall was used to mould the PDMS. After the PDMS was poured into the template, it was left in an oven at 40 ^°^C overnight. This ensured that the PDMS hardened optimally.

Once the PDMS hardened, it was peeled off the template. A small indent is formed with a scalpel in the ends of the middle of the PDMS microfluidic device. This allows two small gold foil strips (Sigma Aldrich 99.99% purity gold) of half a centimetre wide and 5 centimeter length to be placed through the device at each end. The foils almost touch in the middle of the microfluidic device with a small gap. The other end of the foils are perpendicular to the microfluidic devices which allows for clips to be attached to supply an electric current.

Another mixture of PDMS, this time with a 5:2 ratio (to cure faster), was used to glue the PDMS to a glass cover slip. The coverslip glued to the PDMS device is then put in an oven at 80 to 90 ^°^C for roughly 12 to 15 minutes. This ensures that the bacteria can flow through the device.

### C. Fluorophores

The fluorophores used were Thioflavin T (ThT) (Sigma-Aldrich), Propidium Iodide (PI), Syto9, Sytox Green (Thermo Fisher Scientific) and Cy3 (Jena Bioscience). Each fluorophore has a slightly different protocol to ensure a successful experiments.

#### 1. ThT

ThT [58] can serve as a voltage indicator which accumulates inside bacterial cells because of their negative potentials [14] [22] [59]. Our group has shown that this dye was useful to observe the dynamics of light stress in *E. coli, P. aeruginosa* and *B. subtilis* [14].

ThT was used with a final concentration of 10 µM [6] [14] [7] [60]. Furthermore, we have previously shown that the concentration does not affect the growth rate of *DH*5*α* [7].

Each day a fresh batch of ThT liquid solution was prepared with autoclaved water and filter sterilised before usage.

#### 2. Propidium iodide

PI was chosen since it is commonly used to distinguish between live and dead cells. PI is a fluorescent dye that selectively stains dead or permeabilized cells. It is impermeable to the membranes of viable cells but can penetrate damaged cell membranes, binding to nucleic acids, such as DNA, and emitting a red fluorescence upon binding. PI binds to DNA via intercalation, which means that it is inserted between the base pairs of DNA molecules.

This fluorophore was used with a final concentration of 10 µM as per protocol with live-dead assays (Thermo Fisher Scientific). A stock solution was prepared with PI and used for each experiment.

#### 3. Syto9

Syto9 was chosen since it is a standard component with PI in live-dead assays. Syto9 is a membrane-permeable nucleic acid stain that binds to both live and dead cells.

This fluorophore was used with a final concentration of 10 µM as per protocol with live-dead assays. As Syto9 is already a high concentration solution, it was simply inoculated into the liquid bacterial culture with the desired concentration.

#### 4. Sytox Green

Sytox Green was chosen due to similar reasons as PI. Sytox Green is impermeable to live cells with intact membranes but can penetrate the membranes of dead or permeabilized cells. It binds to nucleic acids, such as DNA, and exhibits strong green fluorescence upon binding. It binds to cells’ DNA by intercalation of the base pairs, similar to ThT and Syto9.

This fluorophore was used with a final concentration of 5 µM as per protocol with live-dead assays. As Sytox Green is already a high concentration solution, it was simply inoculated into the liquid bacterial culture with the desired concentration.

#### 5. Cy3

Cy3 was chosen since its emission wavelength is far from ThT and for its ability to covalently label DNA without being disrupted by the environment. When Cy3 is attached to the 5 end of a DNA duplex, the dye stacks on the terminal base pair and stabilizes the duplex [61].

After the PCR product was purified, the DNA labelled with Cy3 was used in a 1:100 ratio of LB. This was later flowed through the microfluidic device which contained the biofilm using a pipette.

### D. Beads

Beads were added to the microfluidic chamber after dilution in a mixture of LB and ThT by adding a total volume of 25 µL, regardless of the beads size.

The smaller beads required a larger pinhole size in the confocal microscope for their fluorescence to be significant. For the 100 nm sulfate beads, a pinhole size of around 80 µm was used, whereas for 30 nm carboxylate beads the pinhole size had to be 100 µm. ThT was used alongside the beads to visualise the cells clearly.

### E. DNA

DNA was obtained from a Lambda DNA product in a “HighFidelity Cy3 PCR Labeling Kit” (Jena Bioscience). It was then purified using a “PCR Purification Kit” (MinElute) and the final product was stored in -20 ^°^C.

### F. Cell Culture and Growth

An agar Lysogeny-Broth (LB) was used to grow single colonies of *E. coli DH5α* from a glycerol stock. When the bacteria successfully grew on the agar plate, a colony was chosen and transferred to a liquid 10 mL LB broth (or 9 mL DM with 1 mL CAA) containing a glass tube. The bacteria then grew overnight in an incubator at 37 ^°^C shaking at 180-250 rpm. We prepared 1L of LB with 10 g sodium chloride (*NaCl*), 10 g tryptone and 5 g of yeast extract. We prepared 1L of Davis minimal medium (DM) as 5.34 g of potassium phosphate (dibasic) *K*_2_*HPO*_4_, 2 g of potassium phosphate (monobasic) *KH*_2_*PO*_4_, 1 g of ammonium sulfate (*NH*_4_)_2_*SO*_4_ and 0.5 g of sodium citrate (trisodium, dihydrate) *Na*_3_*C*_6_*H*_5_*O*_7_(*H*_2_*O*)_2_. After autoclaving we added 1 mL solutions of 10% (w/v) magnesium sulfate *MgSO*_4_ (separately filter sterilised) and 0.2% (w/v) Thiamine (filter sterilized) were added to the stock of DM. Afterwards, 9 mL of this DM stock solution was added and 1 mL of a 2% (w/v) concentration stock solution of CAA was added (i.e. final concentration for CAA is 0.2% (w/v)). Finally, 250 mg per L of glucose was added to the 10 mL solution.

The next day, 100 µl of the liquid culture was added to a fresh 10 ml glass tube containing LB. This is then left until the exponential growth (midlog) phase, attained using a growth curve of the optical density (OD) with time, measured using a spectrometer (JENWAY, Cole-Parmer UK). For DH5*α* the midlog phase was 3 hours and 30 minutes for *OD*_600_ = 0.8.

To sustain the growth temperature at 37 ^°^C, a Perspex microscope chamber heated using an Air-THERM ATX (World Precision Instrument Ltd). The chamber was heated for one hour before the microfluidic device with the bacteria was introduced.

Before inoculating the bacteria into the microfluidic device, Corning ^®^ Cell-tak was added to it and left for around 30 minutes. Then, 300 µL of cells in the midlog phase were stained with the respective fluorophore and added to the Cell-tak filled microfluidic device, which helps the cells adhere to the surface. This was left for another 30 minutes to ensure that the cells were properly stuck to the surface of the microfluidic device. Then the inoculated microfluidic device was taken to the microscope to measure the fluorescence.

#### 1. Biofilm Growth

The biofilm was grown in the microfluidic devices using a media pump (World Precision Instruments UK) which used a syringe filled with LB and ThT. The pump was set for 7 µl per minute and it was left for between 12 to 14 hours. When biofilms were successfully grown then the electric current was applied. The growth temperature was maintained at 37 ^°^C using a Perspex microscope chamber heated using an Air-THERM ATX (World Precision Instrument Ltd.).

### G. Microscopy and Imaging

The microscope was an Olympus IX83 inverted microscope (Klaus Decon Vision) using Blue Lumencor LED excitation (illumination) with 60x (NA 1.42 Plan Apo N) or 100x (NA 1.40 Uplan S Apo) oil immersion objectives. Different filters were used with different flurophores; a CFP filter with ThT (an excitation peak of 440 nm), a mCherry filter with PI (with an excitation peak of 575 nm) and a FITC filter with Syto9 or Sytox Green (an excitation peak of 470 nm).

Images were also collected with a Leica TCS SP8 AOBS inverted gSTED confocal microscope using a 100x HC PL APO (Oil ; STED WHITE). The confocal settings were as follows: pinhole (varies as specified on each experiment), scan speed (400-600 Hz unidirectional) and image size (512 × 512). The supercontinuum laser (8 lines from 470 nm to 670 nm and max power of *<*500 mW) and blue diode (max power of 50 mW) were adjusted to the emission and excitation spectrums of the dyes. For ThT, the excitation was chosen to be at 405 nm (using the blue diode) and a detection range of around 420 - 500 nm. For Cy3, the excitation was chosen at 520 nm and detection range was from 560 to 700 nm. The beads’ ranges were adjusted depending on the peak brightness of the beads as these did not have a specific emission/excitation spectra.

Images were taken every second for either 60 or 120 seconds. Image acquisition was performed using the MetaMorph software (Molecular devices) for the widefield microscope and LAS X v1.8.0.13370 (Leica) for the confocal microscope.

The exposure time was taken at values ranging from 30 to 100 ms depending on the dye used. 30 ms was used for Syto9 and Sytox Green, and 100 ms for PI and ThT. This was due to Syto9 and Sytox green fluorescing brighter than the other dyes.

Several regions of interest, ranging from 5 to 40 regions (i.e. different sections of the sample depending on the microscope’s field of view), taken during the same experimental day, were used to average the data.

### H. Image and Data Analysis

ImageJ (National Institute of Health), MATLAB and Python were used for image analysis. Data analyses, plots (and normalisation) and curve fits were done with MATLAB. The simulation of AC electro-osmosis was done in Python.

Background subtraction in ImageJ was used with a rolling ball radius of 11 µm. The images were also put through a filter in ImageJ depending on the dye used. A median filter of 2.0 was used. The brightness and contrast of the image was regulated to optimize the variation of the intensity seen by eye (this does not affect the data quantitatively).

### I. Electrical Stimulation

The values chosen for the electric fields were first taken from Stratford et al [22]. However, after some experimental optimization, the AC stimulation voltage was taken to be 10 V pp and the frequency 1 kHz to create the largest response of the fluorophores. The AC pulses were 5 seconds long.

## V. CONCLUSIONS

Electrophysiology experiments are important to a wide range of research in medicine and biotechnology. Specifically, the response of excitable matter to an external electrical field is often used to control action potentials that develop across the membranes of cells. We have shown that it is vital to consider the effects of AC electro-osmosis on the microfluidics of samples in such experiments, since they may mask the electrophysiological phenomena demonstrated by excitable matter. It is recommended that when using exogenous flurophores to stain cells, AC electro-osmosis should be considered as an explanation for cellular responses e.g. it causes phantom action potentials during electrical stimulation of *E. coli* suspensions and biofilms.

Furthermore, AC electro-osmosis can be used to drive molecules and nanoparticles through *E. coli* biofilms. This could be relevant to increasing the transport of particles during antibiotic delivery. Small molecule flurophores, DNA and colloidal spheres could all be driven through *E. coli* biofilms using AC electro-osmosis.

## ACKNOWLEDGMENTS

We thank Danna Gifford, Chris Knight and Rowan Green for useful discussions.

We also thank Rebecca Palmer for her help and guidance with the PCR technique and DNA productions.

The Bioimaging Facility microscopes used in this study were purchased with grants from BBSRC, Wellcome and the University of Manchester Strategic Fund. Special thanks goes to Peter March, James Bagnall and Steven Marsden for their help with the microscopy.

## References

[1] M. B. Millerand B. L. Bassler, Quorum sensing in bacteria, Annual Review of Microbiology 55, 165 (2001).

[2] I. Olsen, Biofilm-specific antibiotic tolerance and resistance, European Journal of Clinical Microbiology & Infectious Diseases 34, 877 (2015).

[3] L. K. Vestby, T. Gronseth, R. Simm, and L. L. Nesse, Bacterial biofilm and its role in the pathogenesis of disease, Antibiotics-Basel 9 (2020).

[4] K. J. Loceyand J. T. Lennon, Scaling laws predict global microbial diversity, Proceedings of the National Academy of Sciences of the United States of America 113, 5970 (2016).

[5] T. A. Waigh, The physics of bacteria: from cells to biofilms (Cambridge University Press, 2024).

[6] A. Prindle, J. Liu, M. Asally, S. Ly, J. Garcia-Ojalvo, and G. M. Sueel, Ion channels enable electrical communication in bacterial communities, Nature 527, 59 (2015).

[7] E. Akabuogu, V. Carneiro Da Cunha Martorelli, R. Krašovec, I. Roberts, and T. Waigh, Emergence of ion-channel mediated electrical oscillations in escherichia coli biofilms., eLife (2023).

[8] J. Keenerand J. Sneyd, Mathematical Physiology (Springer New York, NY, 2009).

[9] W. Gerstner, W. M. Kistler, R. Naud, and L. Paninski, Neuronal dynamics: from single neurons to networks and models of cognition (Cambridge University Press, 2014).

[10] E. Skates, H. Delattre, Z. Schofield, M. Asally, and O. S. Soyer, Thioflavin T indicates mitochondrial membrane potential in mammalian cells, Biophysical Reports 3 (2023).

[11] J. Hüserand L. Blatter, Fluctuations in mitochondrial membrane potential caused by repetitive gating of the permeability transition pore, Biochemical Journal 343, 311 (1999).

[12] M. Schwarzlaender, D. C. Logan, I. G. Johnston, N. S. Jones, A. J. Meyer, M. D. Fricker, and L. J. Sweetlove, Pulsing of membrane potential in individual mitochondria: A stress-induced mechanism to regulate respiratory bioenergetics in Arabidopsis, Plant Cell 24, 1188 (2012).

[13] M. Hennes, N. Bender, T. Cronenberg, A. Welker, and B. Maier, Collective polarization dynamics in bacterial colonies signify the occurrence of distinct subpopulations, PLOS Biology 21, 1 (2023).

[14] J. Blee, I. Roberts, and T. Waigh, Membrane potentials, oxidative stress and the dispersal response of bacterial biofilms to 405 nm light, Physical Biology 17, 3 (2020).

[15] J. M. Kralj, D. R. Hochbaum, A. D. Douglass, and A. E. Cohen, Electrical spiking in Escherichia coli probed with a fluorescent voltage-indicating protein, Science 333, 345 (2011).

[16] X. Jin, X. Zhang, X. Ding, T. Tian, C.-K. Tseng, X. Luo, X. Chen, C.-J. Jo, M. C. Leake, and F. Bai, Sensitive bacterial Vm sensors revealed the excitability of bacterial Vm and its role in antibiotic tolerance, Proceedings of the National Academy of Sciences of the United States of America 120 (2023).

[17] A. Hodgkinand A. Huxley, A quantitative description of membrane current and its application to conduction and excitation in nerve, Journal of Physiology-London 117, 500 (1952).

[18] A. Goldberger, L. Amaral, L. Glass, J. Hausdorff, P. Ivanov, R. Mark, J. Mietus, G. Moody, C. Peng, and H. Stanley, Physiobank, physiotoolkit, and physionet - components of a new research resource for complex physiologic signals, Circulation 101, E215 (2000).

[19] J. Heijman, N. Voigt, S. Nattel, and D. Dobrev, Cellular and molecular electrophysiology of atrial fibrillation initiation, maintenance, and progression, Circulation Research 114, 1483 (2014).

[20] O. Kannand R. Kovacs, Mitochondria and neuronal activity, American Journal of Physiology-Cell Physiology 292, C641 (2007).

[21] A. F. Kintzer, K. L. Thoren, H. J. Sterling, K. C. Dong, G. K. Feld, I. I. Tang, T. T. Zhang, E. R. Williams, J. M. Berger, and B. A. Krantz, The protective antigen component of anthrax toxin forms functional octameric complexes, Journal of Molecular Biology 392, 614 (2009).

[22] J. P. Stratford, C. L. A. Edwards, M. J. Ghanshyam, D. Malyshev, M. A. Delise, Y. Hayashi, and M. Asally, Electrically induced bacterial membrane-potential dynamics correspond to cellular proliferation capacity, Proceedings of the National Academy of Sciences of the United States of America 116, 9552 (2019).

[23] P. Marszalek, D. Liu, and T. Tsong, Schwan equation and transmembrane potential induced by alternating electric-field, Biophysical Journal 58, 1053 (1990).

[24] H. P. Schwan, Biophysics of the interaction of electromagnetic energy with cells and membranes, in Biological Effects and Dosimetry of Nonionizing Radiation: Radiofrequency and Microwave Energies, edited by M. Grandolfo, S. M. Michaelson, and A. Rindi (Springer US, Boston, MA, 1983) pp. 213–231.

[25] B. J. Kirby, Micro- and Nanoscale Fluid Mechanics: Transport in Microfluidic Devices (Cambridge University Press, 2010).

[26] H.-C. Changand L. Y. Yeo, Electrokinetically-Driven Microfluidics and Nanofluidics (Cambridge University Press, 2009).

[27] A. Castellanos, A. Ramos, A. González, N. Green, and H. Morgan, Electrohydrodynamics and dielectrophoresis in microsystems:: scaling laws, Journal of Physics D-Applied Physics 36, 2584 (2003).

[28] A. Ramos, Electrokinetics and Electrohydrodynamics in Microsystems (Springer Vienna, 2011).

[29] F. Rieke, W. Bialek, R. de Ruyter van Steveninck, and D. Warland, Spikes : exploring the neural code, Computational neuroscience series. (MIT, Cambridge, Mass. ;, 1999).

[30] E. M. Izhikevich, Dynamical systems in neuroscience : the geometry of excitability and bursting, Computational neuroscience (MIT Press, Cambridge, Mass. ;, 2007).

[31] I. Ershler, A. Pribush, P. Kuzmin, I. Abidor, and M. Yarovaya, A study of electrical breakdown of cell-membrane by the electrooptical method, Biologicheskie Membrany 8, 1327 (1991).

[32] P. N. Tawakoli, A. Al-Ahmad, W. Hoth-Hannig, M. Hannig, and C. Hannig, Comparison of different live/dead stainings for detection and quantification of adherent microorganisms in the initial oral biofilm, Clinical Oral Investigations 17, 841 (2013).

[33] E. Zand, F. Schottroff, C. Schoenher, K. S. Zimmermann, M. Zunabovic-Pichler, and H. Jaeger, Single-staining flow cytometry approach using SYTOX Green to describe electroporation effects on Escherichia coli, Food Control 132 (2022).

[34] N. Green, A. Ramos, A. González, H. Morgan, and A. Castellanos, Fluid flow induced by nonuniform ac electric fields in electrolytes on microelectrodes.: Iii.: Observation of streamlines and numerical simulation -: art. no. 026305, Physical Review E 66 (2002).

[35] E. Secchi, G. Savorana, A. Vitale, L. Eberl, R. Stocker, and R. Rusconi, The structural role of bacterial edna in the formation of biofilm streamers, Proceedings of the National Academy of Sciences of the United States of America 119 (2022).

[36] S. Overballe-Petersen, K. Harms, L. A. A. Orlando, J. V. M. Mayar, S. Rasmussen, T. W. Dahl, M. T. Rosing, A. M. Poole, T. Sicheritz-Ponten, S. Brunak, S. Inselmann, J. De Vries, W. Wackernagel, O. G. Pybus, R. Nielsen, P. J. Johnsen, K. M. Nielsen, and E. Willerslev, Bacterial natural transformation by highly fragmented and damaged dna, Proceedings of the National Academy of Sciences of the United States of America 110, 19860 (2013).

[37] S. Finkeland R. Kolter, Dna as a nutrient novel role for bacterial competence gene homologs, Journal of Bacteriology 183, 6288 (2001).

[38] J.-R. Du, Y.-J. Juang, J.-T. Wu, and H.-H. Wei, Long-range and superfast trapping of dna molecules in an ac electrokinetic funnel, Biomicrofluidics 2, 10.1063/1.3037326 (2008).

[39] E. W. Lee, J. Gehl, and S. T. Kee, Introduction to electroporation, in Clinical Aspects of Electroporation, edited by S. Kee, J. Gehl, and E. Lee (2011) pp. 3–7.

[40] C. F. Munoz, L. de Jaeger, M. H. J. Sturme, K. Y. F. Lip, J. W. J. Olijslager, J. Springer, E. J. H. Wolbert, D. E. Martens, G. Eggink, R. A. Weusthuis, and R. H. Wijffels, Improved dna/protein delivery in microalgae - a simple and reliable method for the prediction of optimal electroporation settings, Algal Research-Biomass Biofuels And Bioproducts 33, 448 (2018).

[41] J. Surre, C. Saint-Ruf, V. Collin, S. Orenga, M. Ramjeet, and I. Matic, Strong increase in the autofluorescence of cells signals struggle for survival, Scientific Reports 8 (2018).

[42] N. Ohtaand T. Takemura, External electric-field effects on fluorescence intensity and decay of pyrimidine vapor, Chemical Physics Letters 169, 611 (1990).

[43] M. Drobizhev, J. N. Scott, P. R. Callis, and A. Rebane, All-optical sensing of the components of the internal local electric field in proteins, IEEE Photonics Journal 4, 1996 (2012).

[44] K. Park, Z. Deutsch, J. J. Li, D. Oron, and S. Weiss, Single molecule quantum-confined stark effect measurements of semiconductor nanoparticles at room temperature, ACS Nano 6, 10013 (2012).

[45] M. Hilczer, S. Traytak, and M. Tachiya, Electric field effects on fluorescence quenching due to electron transfer, 115, 11249 (2001).

[46] S. A. Meredith, Y. Kusunoki, S. D. Connell, K. Morigaki, S. D. Evans, and P. G. Adams, Self-quenching behavior of a fluorescent probe incorporated within lipid membranes explored using electrophoresis and fluorescence lifetime imaging microscopy, Journal of Physical Chemistry B, 1715 (2023).

[47] S. Rani, B. Pitts, and P. Stewart, Rapid diffusion of fluorescent tracers into Staphylococcus epidermidis biofilms visualized by time lapse microscopy, Antimicrobial Agents and Chemotherapy 49, 728 (2005).

[48] L. Cohen, R. Keynes, and B. Hille, Light scattering and birefringence changes during nerve activity, Nature 218, 438 (1968).

[49] W. Ross, B. Salzberg, L. Cohen, A. Grinvald, H. Davila, A. Waggoner, and C. Wang, Changes in absorption, fluorescence, dichroism, and birefringence in stained giantaxons - optical measurement of membrane-potential, Journal of Membrane Biology 33, 141 (1977).

[50] Y. Yang, X.-W. Liu, H. Wang, H. Yu, Y. Guan, S. Wang, and N. Tao, Imaging action potential in single mammalian neurons by tracking the accompanying sub-nanometer mechanical motion, ACS NANO 12, 4186 (2018).

[51] S. Batabyal, S. Satpathy, L. Bui, Y.-T. Kim, S. Mohanty, R. Bachoo, and D. P. Dave, Label-free optical detection of action potential in mammalian neurons, Biomedical Optics Express 8, 3700 (2017).

[52] G. Halnes, T. V. Ness, S. Næss, E. Hagen, K. H. Pettersen, and G. T. Einevoll, Electric Brain Signals: Foundations and Applications of Biophysical Modeling (Cambridge University Press, 2024).

[53] E. Masi, M. Ciszak, L. Santopolo, A. Frascella, L. Giovannetti, E. Marchi, C. Viti, and S. Mancuso, Electrical spiking in bacterial biofilms, Journal of the Royal Society Interface 12 (2015).

[54] X. Zhang, J. Hansing, R. R. Netz, and J. E. DeRouchey, Particle transport through hydrogels is charge asymmetric, Biophysical Journal 108, 530 (2015).

[55] L. M. Rooney, W. B. Amos, P. A. Hoskisson, and G. McConnell, Intra-colony channels in E. coli function as a nutrient uptake system, Nature 14, 2461 (2020).

[56] C. Huang, S. Peretti, and J. Bryers, Plasmid retention and gene-expression in suspended and biofilm cultures of recombinant escherichia-coli dh5 alpha(pmjr1750), Biotechnology and Bioengineering 41, 211 (1993).

[57] S. Soleimani, B. Ormeci, and O. B. Isgor, Growth and characterization of escherichia coli dh5 alpha biofilm on concrete surfaces as a protective layer against microbiologically influenced concrete deterioration (micd), Applied Microbiology and Biotechnology 97, 1093 (2013).

[58] J. Plasekand K. Sigler, Slow fluorescent indicators of membrane potential: A survey of different approaches to probe response analysis, Journal Of Photochemistry And Photobiology B-Biology 33, 101 (1996).

[59] J. Humphries, L. Xiong, J. Liu, A. Prindle, F. Yuan, H. A. Arjes, L. Tsimring, and G. M. Suel, Species-independent attraction to biofilms through electrical signaling, Cell 168, 200 (2017).

[60] J. Liu, R. Martinez-Corral, A. Prindle, D.-Y. D. Lee, J. Larkin, M. Gabalda-Sagarra, J. Garcia-Ojalvo, and G. M. Suel, Coupling between distant biofilms and emergence of nutrient time-sharing, Science 356, 638 (2017).

[61] B. G. Moreira, Y. You, and R. Owczarzy, Cy3 and cy5 dyes attached to oligonucleotide terminus stabilize dna duplexes: Predictive thermodynamic model, Biophysical Chemistry 198, 36 (2015).

