## Supplementary Information for "AC electro-osmosis in bacterial biofilms: a cautionary tale for electrophysiology experiments"

(Dated: October 31, 2024)

### I. SUPPLEMENTARY VIDEOS

The supplementary videos show the phenomena observed when applying a 10 V peak-to-peak (pp) at 10, 30, 50, 70, 90 and 110 seconds in a two minute interval. The electrodes are placed spatially at the top and bottom of the videos unless stated otherwise. The scale used was either a 10x, 60x or 100x magnification (see the Methods section). The videos are all at 30 frames per second.

Videos 1, 2 and 3 show results where the cells do not move and there is a fluorescence dip, which is the main result presented in the article. Videos 4, 5, 6 and 7 show additional evidence for electro-osmotic pumping; that is, the movement of bacterial cells towards one of the electrodes when the voltage pulse is applied. This result appeared whenever the cells did not adhere to the glass surface.

Videos 8 and 9 show how the decrease in fluorophore brightness is not a cellular phenomenon, but instead is a physical artefact caused by AC electro-osmosis of the fluorophores ThT and Sytox Green respectively. There are no cells present and yet the fluorescent dips are still present. The ”subtract background” feature of ImageJ Fiji was not used for these final two videos to highlight the overall brightness as there were no cells present.

### II. SUPPLEMENTARY FIGURES

#### A. Graphs

Figures II.1 to II.6 are from ThT AC electrical stimulation measurements without any cells. The measurements were taken for different frequencies and voltages and plotted against the intensity dip size.

* Biological Physics, Department of Physics and Astronomy, University of Manchester, Oxford Rd., Manchester, M13 9PL, UK.; Division of Infection, Lydia Becker Institute of Immunology and Inflammation, School of Biological Sciences, University of Manchester, Oxford Rd., Manchester, M13 9PT, UK.

† Division of Evolution, Infection and Genomics, School of Biological Sciences, Faculty of Biology, Medicine and Health University of Manchester, Manchester, M13 9PT, UK.

‡ Division of Infection, Lydia Becker Institute of Immunology and Inflammation, School of Biological Sciences, University of Manchester, Oxford Rd., Manchester, M13 9PT, UK.;

§ Biological Physics, Department of Physics and Astronomy, University of Manchester, Oxford Rd., Manchester, M13 9PL, UK.; Photon Science Institute, Alan Turing Building, Oxford Rd, Manchester M13 9PY, UK;

The purpose of this experiment was to determine the relation between the dip size and the voltage.

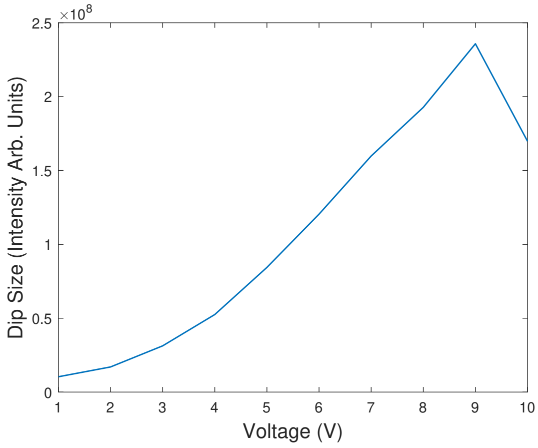


FIG. II.1. ThT fluorescence intensity measurements at different voltages. Analysis of the dip size compared to the voltage supplied in one pulse for 5 seconds. The frequency was kept constant at 100 Hz. The power law factor $b$ in the form $f(x) = ax^b$ is given by 1.298 with 95% confidence interval in the range 0.728 to 1.868. $R^2$ value of 0.8916

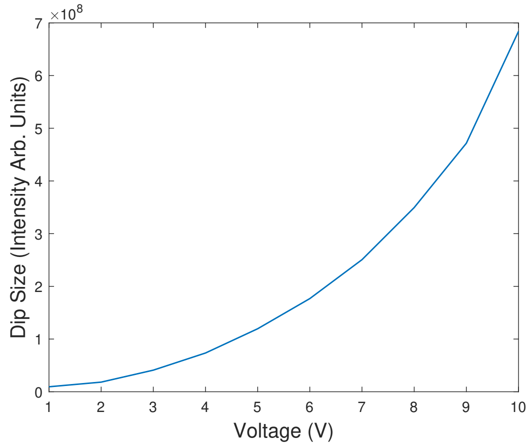


FIG. II.2. ThT fluorescence intensity measurements at different voltages. Analysis of the dip size compared to the voltage supplied in one pulse for 5 seconds. The frequency was kept constant at 100 Hz. The power law factor $b$ in the form $f(x) = ax^b$ is given by 2.655 with 95% confidence interval in the range of 2.387 to 2.923. $R^2$ value of 0.9945.

The frequency dependence was inconclusive from this experiment. However the voltage ($V$) dependency scaled according to equation (I.3) for the dip size ($d$). A power law scaling of $d \sim V^\alpha$ for the dip sizes ($d$) of around $\alpha = 2$ was observed for every frequency.

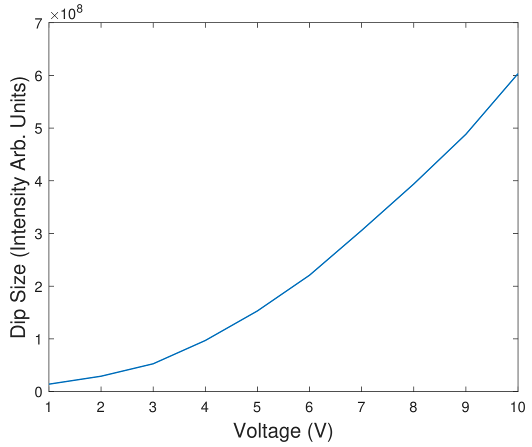


FIG. II.3. ThT fluorescence intensity measurements at different voltages. Analysis of the dip size compared to the voltage supplied in one pulse for 5 seconds. The frequency was kept constant at 100 Hz. The power law factor $b$ in the form $f(x) = ax^b$ is given by 1.965 with 95% confidence interval in the range 1.912 to 2.018. $R^2$ value of 0.9996.

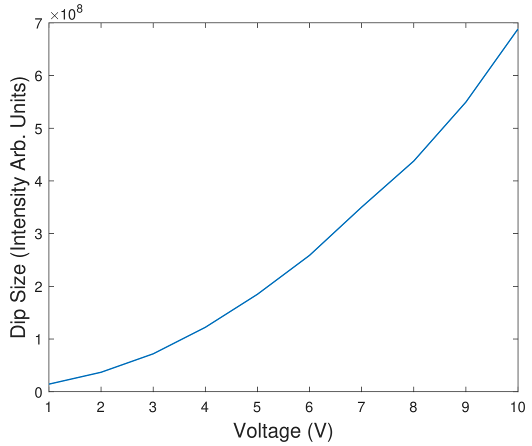


FIG. II.4. ThT fluorescence intensity measurements at different voltages. Analysis of the dip size compared to the voltage supplied in one pulse for 5 seconds. The frequency was kept constant at 100 Hz. The power law factor $b$ in the form $f(x) = ax^b$ is given by 1.888 with 95% confidence interval in the range to 1.829,1.947. $R^2$ value of 0.9994.

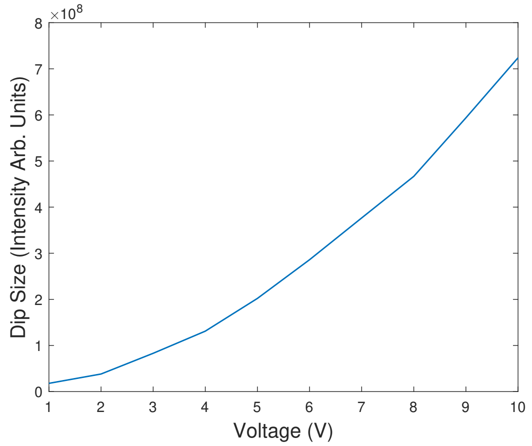


FIG. II.5. ThT fluorescence intensity measurements at different voltages. Analysis of the dip size compared to the voltage supplied in one pulse for 5 seconds. The frequency was kept constant at 100 Hz. The power law factor $b$ in the form $f(x) = ax^b$ is given by 1.834 with 95% confidence interval in the range 1.783 to 1.884. $R^2$ value of 0.9995.

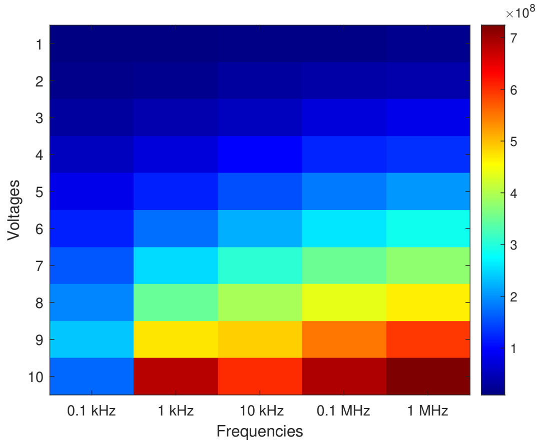


FIG. II.6. ThT fluorescence intensity measurements at different applied AC voltages. Colourmap of the dip size as a function of the voltage. This graph shows the data from Figure II.1 to Figure II.5.

#### B. Microfluidic Device Template

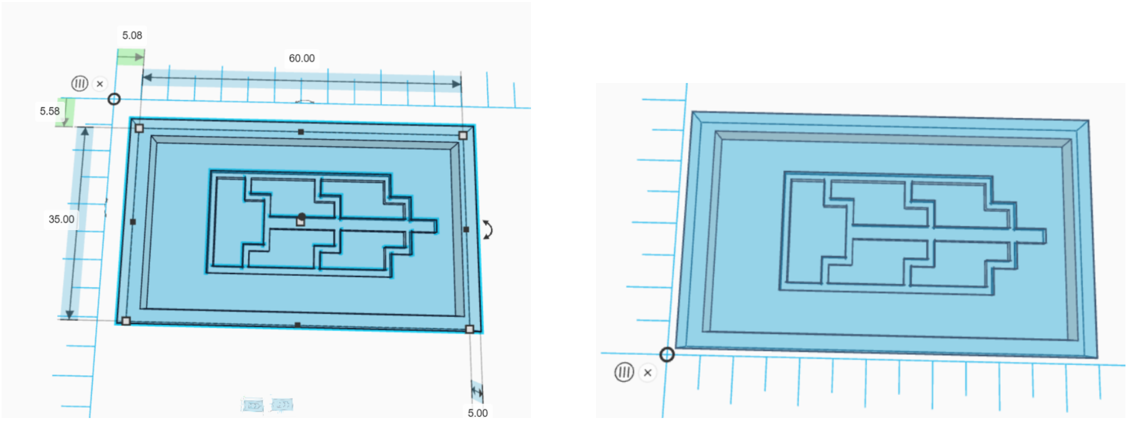


(a) (b)

FIG. II.7. Template of 3D microfluidic device. (a) Shows the template's dimensions. It is 35 mm wide, 60 mm long and 5 mm deep. The indents where the bacteria will sit are 2 mm deep. (b) Shows the template without any ruler for ease of view.

#### C. Simulation Details

To simulate AC electro-osmosis around two parallel electrodes using Matlab, coupled electric and fluid equations (Poisson-Nernst-Planck and Navier-Stokes) were solved numerically. The solutions were created using the finite element method.

The geometry was defined via the length, width and spacing of the electrodes. The electrode spacing was the most important factor. Electrical properties were defined: voltage, frequency and phase were defined for our experimental set up and Stratford et al's. Fluid properties were defined via the dynamic viscosity ($\mu$), density ($\rho$) and permittivity ($\epsilon$).

Laplace's equation (equation II.1) was used to predict the electric potential distribution,

$$\nabla^2 \phi = 0, \tag{II.1}$$

where $\nabla^2 \phi$ is the Laplacian of the scalar potential $\phi$, which in this case is the electric potential.

Navier-Stokes equations (equation II.2) predicted the fluid flow coupled with Laplace's equation,

$$\rho \left( \frac{\partial \mathbf{u}}{\partial t} + \mathbf{u} \cdot \nabla \mathbf{u} \right) = -\nabla p + \mu \nabla^2 \mathbf{u} + \mathbf{f}, \tag{II.2}$$

where $\rho$ is the density of the fluid, $\mathbf{u}$ is the velocity field, $p$ is the pressure, $\mu$ is the dynamic viscosity, $\mathbf{f}$ represents the external forces, $\nabla\mathbf{u}$ is the gradient of the velocity and $\nabla^2\mathbf{u}$ is the Laplacian of the velocity. In practice, the inertial terms were negligible in the simulations and the results from the Navier-Stokes equation simulations were indistinguishable from Stokes equation simulations.

At the electrode, the electro-osmotic slip velocity $u$ is

$$u = -\frac{\epsilon}{\eta}\Delta\phi_d\frac{\partial\phi}{\partial x} = \frac{\epsilon}{\eta}\Delta\phi_d E_x, \tag{II.3}$$

where $E_x$ is the tangential field just outside the diffuse layer, $\eta$ is the fluid viscosity and $\Delta\phi_d = \phi - \psi$ represents the difference between the potential $\phi$ on the outer side of the diffuse layer and the potential $\psi$ on the inner side of this layer, at the nonslip plane.

The boundary conditions for the electrodes (via the AC signal) and the domain edges were applied during numerical solution. Both Navier-Stokes equations and Laplace's equations were solved using Finite Element Methods (FEMs) in Matlab.

The boundary conditions and simulations followed the work of Green et al [1]. Thus the Debye-Huckel model was used to characterise the voltage and slip velocity on the electrode. On the surface of the electrodes, the boundary condition was

$$\sigma Z_{DL}\frac{\partial\phi}{\partial y} = \phi - V. \tag{II.4}$$

where $Z_{DL}$ is the impedance of the double layer with $Z_{DL} = 1/i\omega C$ with $C = \epsilon/\lambda_D$ and $\lambda_D$ is the Debye length. This equation describes the potential at the Debye length. The other boundary conditions were that at the mirror plane $(x = 0)$ the potential is zero and there is a Neumann boundary condition $\partial\phi/\partial n = 0$ everywhere else.

The velocities also have boundary conditions. At the electrode, the $x$ component is governed by Equation II.3 and the velocity $y$ component is assumed to be zero. At the mirror plane $(x = 0)$, the $x$ component of the velocity is 0 and there is a Neumann boundary condition for the velocity in $y$. $\partial v/\partial n = 0$. $u$ and $v$ are assumed to be zero everywhere else at the boundaries.

The domain was discretized by using a grid to represent the space between and around the electrodes. The electric field was computed by using the potential. The fluid velocity was determined based on the electric field and boundary conditions. Finally, plots of the potential contours, magnitude of velocity and

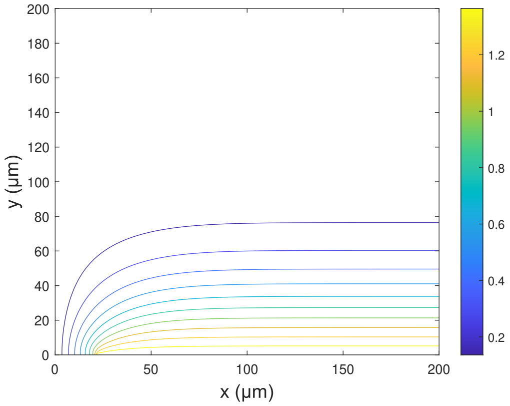


(a)

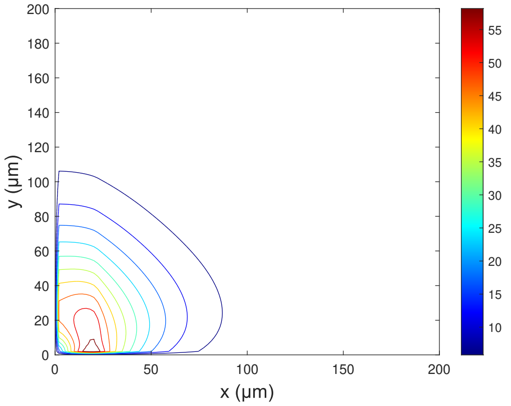


(b)

FIG. II.8. Microfluidic simulation of AC electro-osmosis from two linear electrodes separated by a gap of 50 μm and an AC voltage of 3 V with 100 Hz i.e. Stratford's et al parameters. The view is from the side of one of the two electrodes. A finite element model is solved based on a combination of Stokes equation and the Laplace equation alongside the Helmholtz-Smoluchowski formula. (a) Shows the electrical potential with the colour-bar in Volts and (b) shows the streamline flow contour plot for the fluid mechanics. The colour-bar is in μm/s. There is a plane of mirror symmetry at x = 0 for both (a) and (b)

fluid flow vectors were obtained.

#### D. Other Simulation Geometries

Figure II.8, Stratford et al's geometry [2] shows that although the velocity streamline flow is different to simulations based on our experimental parameters, there is a clear indication that there is a fluid flow in the microfluidic device, which could transport the fluorophores and cause phantom action potentials in the fluorophore intensities.

Figure II.9 shows the streamline velocities for two thick electrodes separated by a gap of 25 μm. This was done to ensure that the simulations agree with the experiments of Green et al [1] i.e. to ensure they were correctly benchmarked.

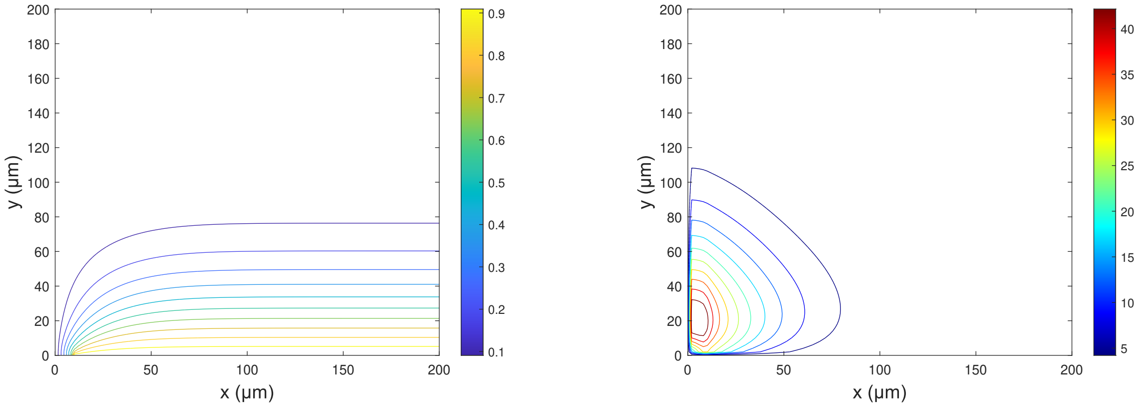


(a) (b)

FIG. II.9. Microfluidic simulation of AC electro-osmosis from two linear electrodes separated by a gap of 25 µm and an AC voltage of 0.5 V with 1000 Hz i.e. Green et al parameters. The view is from the side of one of the two electrodes. A finite element model is solved based on a combination of Stokes equation and the Laplace equation alongside the Helmholtz-Smoluchowski formula. Figure (a) Shows the electrical potential with the colour-bar in volts and (b) shows the streamline flow contour plot for the fluid mechanics. The colour-bar is in µm/s. There is a plane of mirror symmetry at x = 0 for both (a) and (b)

[1] N. Green, A. Ramos, A. González, H. Morgan, and A. Castellanos, Fluid flow induced by nonuniform ac electric fields in electrolytes on microelectrodes.: Iii.: Observation of streamlines and numerical simulation -: art. no. 026305, Physical Review E **66** (2002).

[2] J. P. Stratford, C. L. A. Edwards, M. J. Ghanshyam, D. Malyshev, M. A. Delise, Y. Hayashi, and M. Asally, Electrically induced bacterial membrane-potential dynamics correspond to cellular proliferation capacity, Proceedings of the National Academy of Sciences of the United States of America **116**, 9552 (2019).
